# Structural basis for constitutive activation of oncogenic gp130 mutants in human inflammatory hepatocellular adenomas

**DOI:** 10.1101/2025.11.11.685192

**Authors:** Yi Zhou, Jing Cao, Trudy Ramlall, Luke L. McGoldrick, Jennifer Jones, Annan S. I. Cook, Raymond Leidich, Mark W. Sleeman, William C. Olson, Panayiotis E. Stevis, Matthew C. Franklin

**Author notes:** Correspondence (Y.Z.); (P.E.S.); (M.C.F.). These authors contribute equally (Jing Cao, Trudy Ramlall, Luke L. McGoldrick).

## Abstract

Inflammatory hepatocellular adenomas (IHCAs) are benign liver tumors primarily driven by somatic mutations in the gp130 receptor, resulting in its constitutive activation. To uncover the structural mechanisms underlying this activation, we employed cryo-electron microscopy and mutagenesis to study five representative IHCAs-associated gp130 mutants and two engineered mutants with ligand-independent activity. Most mutants formed “X”-shaped dimers through a 3D domain-swapping mechanism, characterized by large juxtamembrane separations and reliant on D1 domain-mediated “tip-to-tip” interactions for dimer clustering and activation. A rare mutant displayed a distinct “Y”-shaped dimer conformation, with closely aligned juxtamembrane domains enabling direct activation independent of dimer clustering. These findings highlight the diversity of gp130 mutant activation mechanisms and provide a foundation for developing therapeutic strategies targeting aberrant gp130 signaling in IHCAs.

## Introduction

Inflammatory hepatocellular adenomas (IHCAs) are the most prevalent subtype of hepatocellular adenomas (HCAs) and are classified as benign liver tumors^1^. These tumors primarily affect females and are associated with risk factors including oral contraceptive use, obesity, and alcohol abuse^2,3^. IHCAs are distinct in their pathological and molecular characteristics, such as polymorphic inflammatory cell infiltrates, activation of acute-phase proteins, and persistent phosphorylation of signal transducer and activator of transcription 3 (STAT3), a transcription factor that plays a pivotal role in inflammation and tumorigenesis. These attributes highlight the inflammatory nature of IHCAs, which, while typically benign, carry a potential risk of progression into hepatocellular carcinomas (HCCs)^1^.

Previous studies have identified specific genetic mutations that drive the development of IHCAs^1,4,5^. Approximately 5% of these tumors are associated with activating mutations in STAT3. However, the majority of cases (∼60%) are driven by activating somatic mutations in the interleukin-6 signal transducer (IL6ST) gene, which encodes the gp130 receptor^1^. Notably, activating mutations in gp130 are also found in 2% of HCCs^4,6,7^. These findings highlight the oncogenic properties of gp130 mutations that drive constitutive activation^8^.

The gp130 receptor is a critical signaling transducer shared among the interleukin-6 (IL-6) family cytokines, including IL-6, IL-11, Ciliary neurotrophic factor (CNTF), Cardiotrophin-like cytokine factor 1 (CLCF1), Leukemia inhibitory factor (LIF), Oncostatin M (OSM), Cardiotrophin-1 (CT-1), IL-27, IL-35, and IL-39^9^. Most of these cytokines require a secondary signaling receptor, such as the LIF receptor (LIFR), OSM receptor (OSMR), or IL-27 receptor alpha (IL-27Rα), to form heterodimers with gp130 for signal transduction. However, IL-6 and IL-11 uniquely signal through gp130 homodimers. Structurally, the extracellular domains (ECDs) of gp130 consist of an N-terminal immunoglobulin (Ig)-like domain (D1), a cytokine-binding homology region (CHR, D2-D3), and three membrane-proximal fibronectin type III domains (FNIII, D4-D6)^9,10^. gp130 D2 and D3 exhibit a common β-sandwich fold composed of a three-stranded (A, B, and E) and a four-stranded (C, C’, F, and G) β-sheet. The cytokine-binding site is situated in two interstrand loop regions: the EF loop in D2 and the BC loop in D3^9–14^.

IL-6 is a multifunctional cytokine that plays a critical role in processes such as inflammation, tissue regeneration, and tumor development^15^. IL-6 initially binds to its receptor α-subunit IL-6Rα and subsequently interacts with the signal-transducing β-subunit gp130, forming a hexametric signaling complex^11^. In this assembly, the acute bend of gp130 at D4D5 positions the juxtamembrane domains of two gp130 molecules into close proximity, enabling the dimerization and activation of intracellularly bound Janus kinases (JAKs), which further phosphorylate and activate the STAT family proteins, including STAT3 and STAT1^9^.

The constitutive activation of gp130 in IHCAs has been attributed to gain-of-function somatic mutations^1,4^. The spectrum of IHCAs-associated gp130 mutations includes 20 distinct in-frame deletions, one missense substitution, and several cases of in-frame insertions coupled with deletions. These mutations predominantly cluster around the EF loop of domain D2, a region critical for IL-6 binding^1,4^. Two amino acids within this hotspot, Y190 and F191, are crucial for mediating the interaction between gp130 and IL-6, with the majority of in-frame deletions specifically impacting these residues. Two most frequent deletions, S187–Y190 deletion (gp130 ΔSY) and Y186–Y190 deletion (gp130 ΔYY), have been biochemically and functionally characterized by several groups^1,4,13,16^. The gp130 ΔSY mutant was found to homodimerize in the absence of a ligand, and its constitutive activity was inhibited by the overexpression of wild-type (WT) gp130^4^. The gp130 ΔYY mutant also exhibited ligand-independent homodimerization, with the D1 domain being critical for activation but not required for dimerization^13,16^. Notably, an inhibitory effect on gp130 ΔYY constitutive activation was not observed for soluble WT gp130 extracellular domain protein^16^. Moreover, an atypical in-frame deletion of four residues, A418–F421 deletion (gp130 ΔAF), located in the D4D5 linker region, was identified in one tumor. This mutation also led to gp130 dimerization and activation^1^.

However, it is not clear whether other IHCAs-related gp130 mutations, such as the K173-D177 deletion (gp130 ΔKD), E195-V196 deletion (gp130 ΔEV), and P216H mutation (gp130 P216H), can also drive homodimerization. Furthermore, the exact molecular mechanisms governing the dimerization and activation of these gp130 mutants are yet to be fully understood.

In this study, we explored the structural mechanisms driving the constitutive activation of gp130 mutants. Utilizing cryo-electron microscopy (cryo-EM) and mutagenesis, we examined five representative gp130 variants identified in IHCAs alongside two engineered mutants with constitutive activity, uncovering distinct mutant conformations and signaling behaviors. Most mutants were found to rely on 3D domain swapping to form “X”-shaped dimers with large juxtamembrane D6 domain separations, requiring D1-mediated “tip-to-tip” interactions for effective dimer clustering and activation. In contrast, the rare ΔAF mutant displayed a unique “Y”-shaped dimer conformation, characterized by closely aligned juxtamembrane domains that enable direct receptor activation independent of dimer clustering. These results underscore the structural diversity of gp130 mutants and deepen our understanding of the ligand-independent activation mechanisms, offering valuable insights into the intervention of gp130-mediated signaling pathways in IHCAs and related pathologies.

## Results

### 3D domain swapping drives dimerization of the extracellular domain of IHCAs-associated gp130 S187-Y190 deletion mutant

The gp130 S187-Y190 deletion mutant is the most commonly identified gp130 variant in IHCAs^1^. The extracellular domain of this mutant (gp130 ΔSY ECD) was expressed as a mixture of monomeric and dimeric forms, as evidenced by Size Exclusion Chromatography (SEC) profile, SEC-MALS (Multi-Angle Light Scattering) analysis, and Native polyacrylamide gel electrophoresis (PAGE, Supplementary Fig. 1a–d). The dimeric (Pool A) and monomeric (Pool C) proteins were subjected to cryo-EM analysis in the presence of the Fab fragment of a commercial anti-gp130 antibody B-S12, which was used as a fiducial marker to aid particle alignment and structure determination^10^. 2D classification of the Pool A sample showed clear features of a dimeric gp130 form as compared to the WT gp130 ECD sample (Supplementary Fig. 1e, f). No good 2D averages were obtained for the Pool C sample (Supplementary Fig. 1g), suggesting that the monomeric form of the mutant might be misfolded. The Fab fragment of a second anti-gp130 antibody, H4xH16683P2 (designated as 16683), was further added to the Pool A/B-S12 sample, leading to a more uniform particle orientation than when the B-S12 Fab was used alone (Supplementary Fig. 1h). A 3.24 Å resolution cryo-EM map was obtained for this complex, featuring a 2:2:2 stoichiometry with 2-fold symmetry (Supplementary Fig. 2a-e). The 16683 Fab binds to a site distant from the gp130 ΔSY dimer interface, whereas the B-S12 Fab binds to the opposite side of the interface in D2 domain (Supplementary Fig. 2e-f), thereby excluding the possibility that these Fabs contribute to the dimerization of the mutant.

Surprisingly, the two gp130 ΔSY molecules within the dimer adopt a “back-to-back” arrangement, with their cytokine-binding domains (D2D3) leaning against each other, creating an “X”-shaped conformation (Fig. 1a). The dimerization is driven by a 3D domain-swapping mechanism, wherein the β_C’_-C’E loop-β_E_ module within D2 assumes an extended conformation (Fig. 1b) and is exchanged between the two molecules (Fig. 1a, c, d). The two gp130 molecules in the dimer are oriented roughly in the same plane of the dimer interface (Fig. 1c). The swapped β_C’_-C’E loop-β_E_ module adopts a conformation resembling that of WT gp130 and interacts with the counterpart molecule in the dimer, forming a chimeric D2 structure that mirrors the folding of D2 in WT gp130 (Fig. 1d-f).

**Fig. 1.**
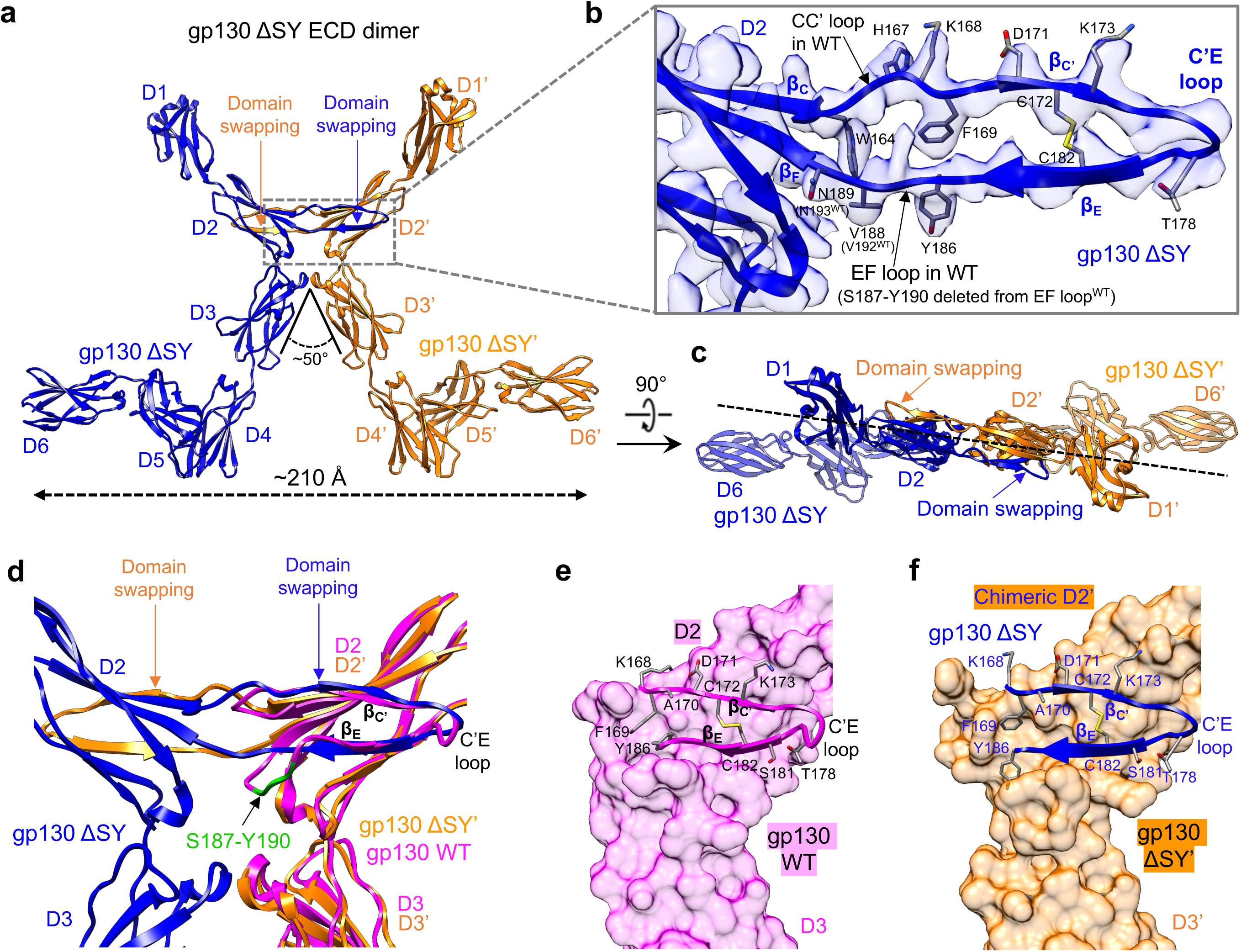
Dimerization of the extracellular domain of the IHCAs-associated gp130 S187-Y190 deletion mutant (gp130 ΔSY ECD) is driven by 3D domain swapping. **a** Side view of cryo-EM structure of the gp130 ΔSY ECD dimer shows 3D domain swapping of a module in D2. The two gp130 ΔSY molecules are colored in blue and orange, respectively. Approximate distance between the two gp130 ΔSY D6 domains is estimated. **b** Zoomed-in view of atomic model in transparent density of D2 of the blue gp130 ΔSY molecule shows an extended conformation of the β_C’_-C’E-β_E_ module. A few residues in the module are shown in stick representation. **b** is in the same orientation as **a**. **c** Top-down view of cryo-EM structure of the gp130 ΔSY ECD dimer. Dash line indicates the plane of the dimer interface. **d** Overlay of gp130 ΔSY ECD dimer structure and WT gp130 structure (PDB: 1PVH). The two gp130 ΔSY molecules are colored in blue and orange, respectively. WT gp130 is colored in magenta. **e** WT gp130 structure (PDB: 1PVH) shown in the same orientation as **d**. The β_C’_-C’E-β_E_ module in D2 is shown in cartoon while the rest of the molecule is shown in surface representation. Residues involved in the packing of the β_C’_-C’E-β_E_ module against the rest of D2 are shown in stick representation. **f** In the gp130 ΔSY mutant dimer, the β_C’_-C’E-β_E_ module from the one molecule (blue gp130 ΔSY shown in cartoon) is packed against the counterpart molecule (orange gp130 ΔSY’ shown in surface representation) to make a chimeric D2 analogous to how D2 is folded in WT gp130.

### Hydrophobic and hydrogen bonding interactions around the EF loop stabilize the folded conformation of the **β**_C’_-C’E-**β**_E_ module in D2 of WT gp130

In WT gp130 D2, the β_C’_-C’E-β_E_module folds back by bending the adjacent CC’ loop and EF loop (Fig. 2a). This folded conformation is likely stabilized by an intricate network of hydrophobic and hydrogen bonding interactions mediated by residues located within and adjacent to the EF loop (Fig. 2a)^13^. Two distinct hydrophobic clusters, V189-V192-I194-P216 and Y190-F191-V252, form hydrophobic cores that flank the EF loop, providing structural support. Additionally, hydrogen bonds formed by D215 with the backbone amides of F191 and V192 in the EF loop may play a crucial role in precisely positioning the loop, ensuring that the β_C’_-C’E-β_E_ module adopts a thermodynamically favorable folded conformation.

**Fig. 2.**
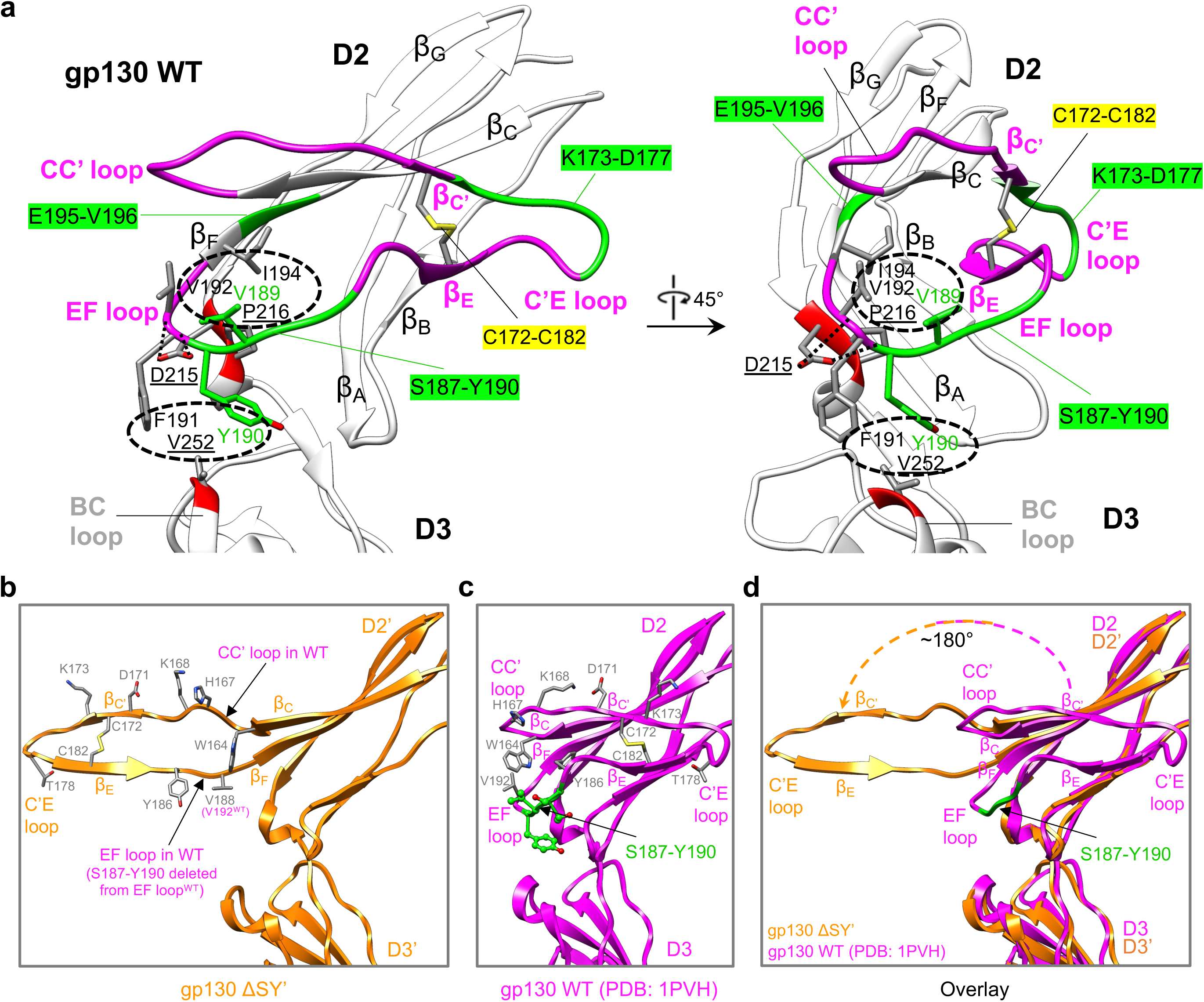
The folded conformation of the β_C’_-C’E-β_E_ module in D2 of WT gp130 is stabilized by hydrophobic and hydrogen bonding interactions around the EF loop, which are disrupted in the S187-Y190 deletion mutant and potentially in other mutants. **a** The structure of WT gp130 D2D3 (PDB: 1PVH) is presented in two orientations, with the β_C’_-C’E-β_E_ module and the adjacent CC’ and EF loops highlighted in magenta. The deleted regions in three constitutively active gp130 mutants (gp130 ΔKD, gp130 ΔSY, and gp130 ΔEV)—K173-D177 in the C’E loop, S187-Y190 in the EF loop, and E195-V196 in β_F_—are highlighted in green. The mutated residues—D215 and P216 in the short helix of D2, and V252 in the BC loop of D3—in another three constitutively active mutants (gp130 D215G, gp130 P216H, and gp130 V252G) are highlighted in red. The EF loop in the D2 domain is stabilized by two hydrophobic cores (V189-V192-I194-P216 and Y190-F191-V252) and two hydrogen bonds formed between D215 and the backbone amides of F191 and V192. The hydrophobic cores are outlined with dashed oval circles, while the hydrogen bonds are depicted using dashed lines. All mentioned residues, along with the two cysteines (C172 and C182) that form the disulfide bond (C172-C182) linking β_C’_ and β_E_, are displayed in stick representation. **b** Zoomed-in view of the atomic model of D2D3 in the orange gp130 ΔSY’ molecule in Fig.1**a** with a few residues in the β_C’_-C’E-β_E_ module being shown in stick representation. **c** WT gp130 structure (PDB: 1PVH) is shown in the same orientation as **b**. The corresponding residues highlighted in **b** are also shown in stick representation. The deleted residues (S187-Y190) in the mutant is colored in green. **d** Overlay of **b** and **c** demonstrates a ∼180° unfolding of the β_C’_-C’E-β_E_ module in the gp130 ΔSY’ mutant (orange) compared to the WT gp130 (magenta).

Notably, the four residues deleted in the gp130 ΔSY mutant, S187–Y190, are situated within the EF loop of gp130. Deletion of these residues will disrupt the two hydrophobic cores, which is sufficient to trigger a ∼180° unfolding of the β_C’_-C’E-β_E_ module (Fig. 2b-d), inducing domain swapping and dimerization (Fig. 1). In line with this, it has been shown that the shortening of hinge loops in other proteins by 1–6 residues can create conformational strain that destabilizes the monomeric structure, promoting the formation of domain-swapped oligomers where the loop adopts an energetically favorable extended conformation^17,18^.

### Mutations D215G and V252G, engineered to disrupt hydrogen bonding and hydrophobic interactions around the EF loop in gp130 D2, induce domain-swapping and dimerization

Two point mutations, D215G and V252G, were previously introduced into gp130 to disrupt the hydrogen bonding and hydrophobic interactions mediated by D215 and V252, respectively, which appear to be critical for proper positioning of the EF loop in gp130 D2 (Fig. 2a). Both mutations resulted in constitutive activation of gp130^13^. However, it remains unclear whether these engineered mutations also induce homodimerization of gp130, similar to the gp130 ΔSY and ΔYY variants observed in IHCAs.

The ECD proteins of the two engineered mutants, D215G and V252G, were purified and characterized using cryo-EM in the presence of B-S12 Fab and 16683 Fab, yielding cryo-EM structures with resolutions of 3.38 Å and 3.82 Å, respectively (Supplementary Fig. 3-5).

Interestingly, like gp130 ΔSY, both mutants exist as homodimers, with the cytokine-binding sites in the D2D3 domains of the two monomers packed against each other to form an “X”-shaped structure (Fig. 3a-f). However, in comparison to the gp130 ΔSY dimer, which features an angle of ∼50° between its two D3 domains (Fig. 1a), the gp130 D215G dimer and V252G dimer exhibit angles of roughly 30° and 90° between D3 domains, respectively. In addition, the gp130 D215G dimer, much like the gp130 ΔSY dimer, exhibits a planar configuration where the two gp130 molecules lie approximately in the same plane of the dimer interface (Fig. 1c, 3c). This contrasts with the gp130 V252G dimer, where the two gp130 molecules are tilted away from the dimer interface plane in opposite directions (Fig. 3f). These angle and planar arrangement differences result in a more “closed” conformation for the D215G dimer and a more “open” conformation for the V252G dimer (Fig. 3a, d). The dimerization of these two engineered mutants is also driven by 3D domain swapping. In both mutant dimers, the β_C’_-C’E loop-β_E_ module in D2 of each monomer adopts an extended, unfolded conformation (Fig. 3b, e), interacting with the opposing D2 rather than folding back against its own D2 (Fig. 3a-f). These data suggest that disrupting hydrogen bonding or hydrophobic interactions critical for the proper positioning of the EF loop within D2—whether through engineered mutations or mutations associated with IHCAs—can serve as a driving force for gp130 domain swapping and dimerization.

**Fig. 3.**
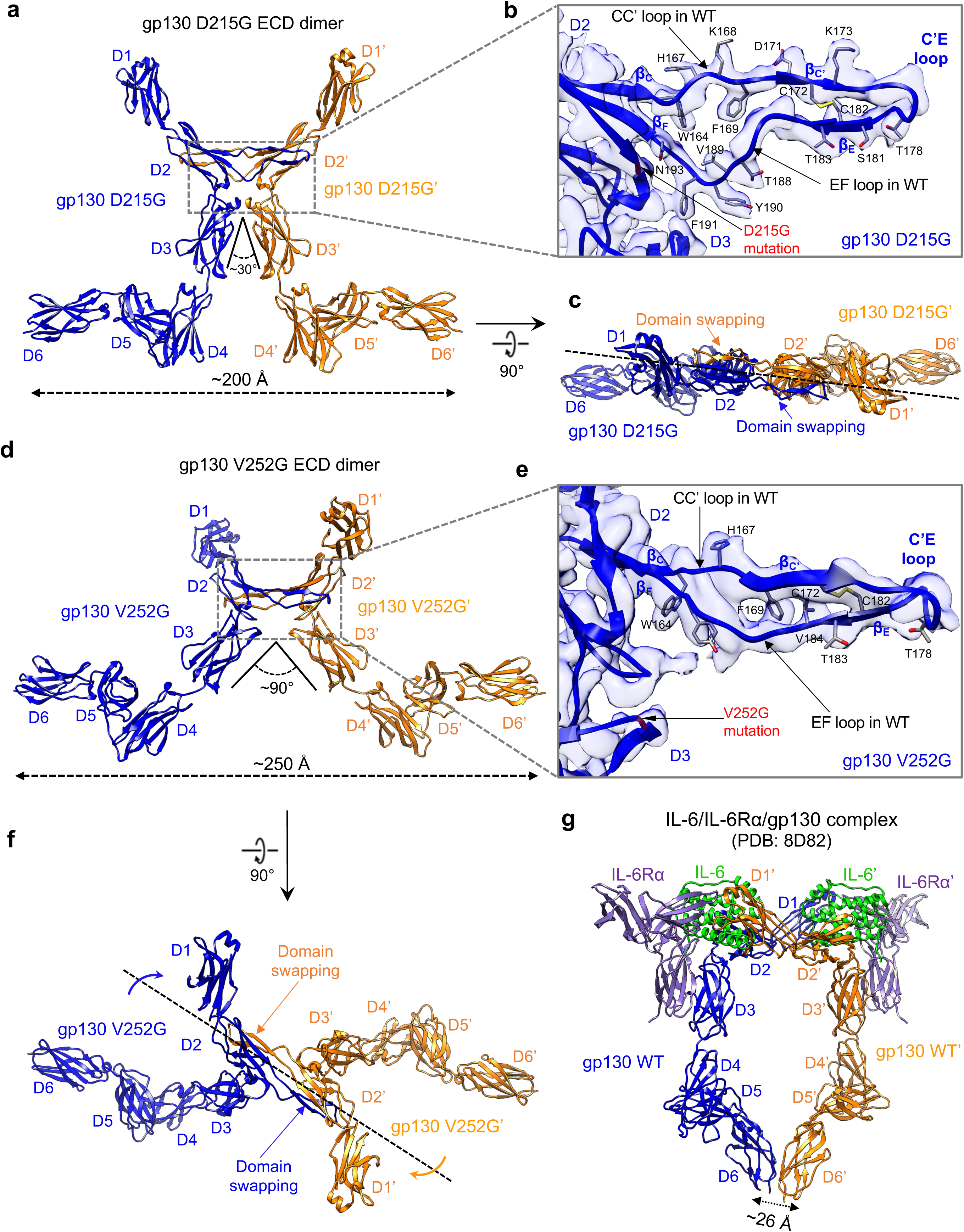
3D domain swapping drives dimerization of the extracellular domain of two engineered constitutively active gp130 mutants (D215G and V252G). **a** Side view of cryo-EM structure of the gp130 D215G ECD dimer shows 3D domain swapping of the β_C’_-C’E-β_E_ module in D2. The two gp130 D215G molecules are colored in blue and orange, respectively. Approximate distance between the two D6 domains is estimated. **b** Zoomed-in view of atomic model in transparent density of D2 of the blue gp130 D215G molecule shows an extended conformation of the β_C’_-C’E-β_E_ module. A few residues in the module are shown in stick representation. **b** is in the same orientation as **a**. **c** Top-down view of cryo-EM structure of the gp130 D215G ECD dimer. Dash line indicates the plane of the dimer interface. **d** Side view of cryo-EM structure of the gp130 V252G ECD dimer shows 3D domain swapping of the β_C’_-C’E-β_E_ module in D2. **e** Zoomed in view of atomic model in transparent density of D2 of the blue gp130 V252G molecule shows an extended conformation of the β_C’_-C’E-β_E_ module. A few residues in the module are shown in stick representation. **e** is in the same orientation as **d**. **f** Top-down view of cryo-EM structure of the gp130 V252G ECD dimer. Dash line indicates the plane of the dimer interface. Arrows indicate the tilting of the two gp130 molecules away from the plane of the dimer interface. **g** Structure of the IL-6/IL-6Rα/gp130 complex (PDB: 8D82). The distance between the two D6 domains is estimated.

### Distinct conformations of constitutively active gp130 mutant dimers compared to WT gp130 homodimer in the IL-6 receptor complex

In the IL-6 receptor complex, two WTgp130 molecules interact with two IL-6/IL-6Rα subcomplexes to adopt an “O”-shaped conformation (Fig. 3g)^9^, which positions the two juxtamembrane D6 domains in close proximity (∼26 Å), a geometry crucial for the initiation of downstream signaling. In contrast, the three constitutively active gp130 mutants, ΔSY, D215G, and V252G, all adopt a distinct “X”-shaped dimer conformation. In this arrangement, the juxtamembrane D6 domains are oriented away from each other, spanning distances of approximately 210 Å, 200 Å, and 250 Å, respectively (Fig. 1a, 3a, 3d), which are too far away to support the dimerization and activation of intracellularly bound JAK kinases^19–22^. Therefore, from a structural standpoint, it is difficult to explain how these gp130 mutants exhibit constitutive activity in cells^4,13^.

### D1 of full-length gp130 **Δ**SY mutant mediated clustering of the mutant dimers via “tip-to-tip” interactions

The gp130 Y186–Y190 deletion mutant (gp130 ΔYY), which has one additional residue deletion compared to gp130 ΔSY, is the second most frequently identified gp130 variant in IHCAs.

Studies have shown that truncation of D1 domain in gp130 ΔYY impairs its constitutive activation^13,16^, highlighting the importance of the D1 domain in activating this mutant. To determine if D1 is similarly essential for gp130 ΔSY activation, a STAT3 luciferase reporter assay was conducted (Fig. 4a). The ligand-independent activity of gp130 ΔSY was greatly reduced following D1 deletion, emphasizing the critical role of D1 in mediating formation of the gp130 ΔSY signaling complex. Notably, unlike the IL-6 signaling complex where gp130 D1 domain directly contributes to complex assembly (Fig. 3g), D1 in gp130 ΔSY mutant neither facilitates dimer formation nor interacts with the rest of the dimer (Fig. 1a), which poses a challenge in understanding the structural basis for the necessity of D1 in mutant activation.

**Fig. 4.**
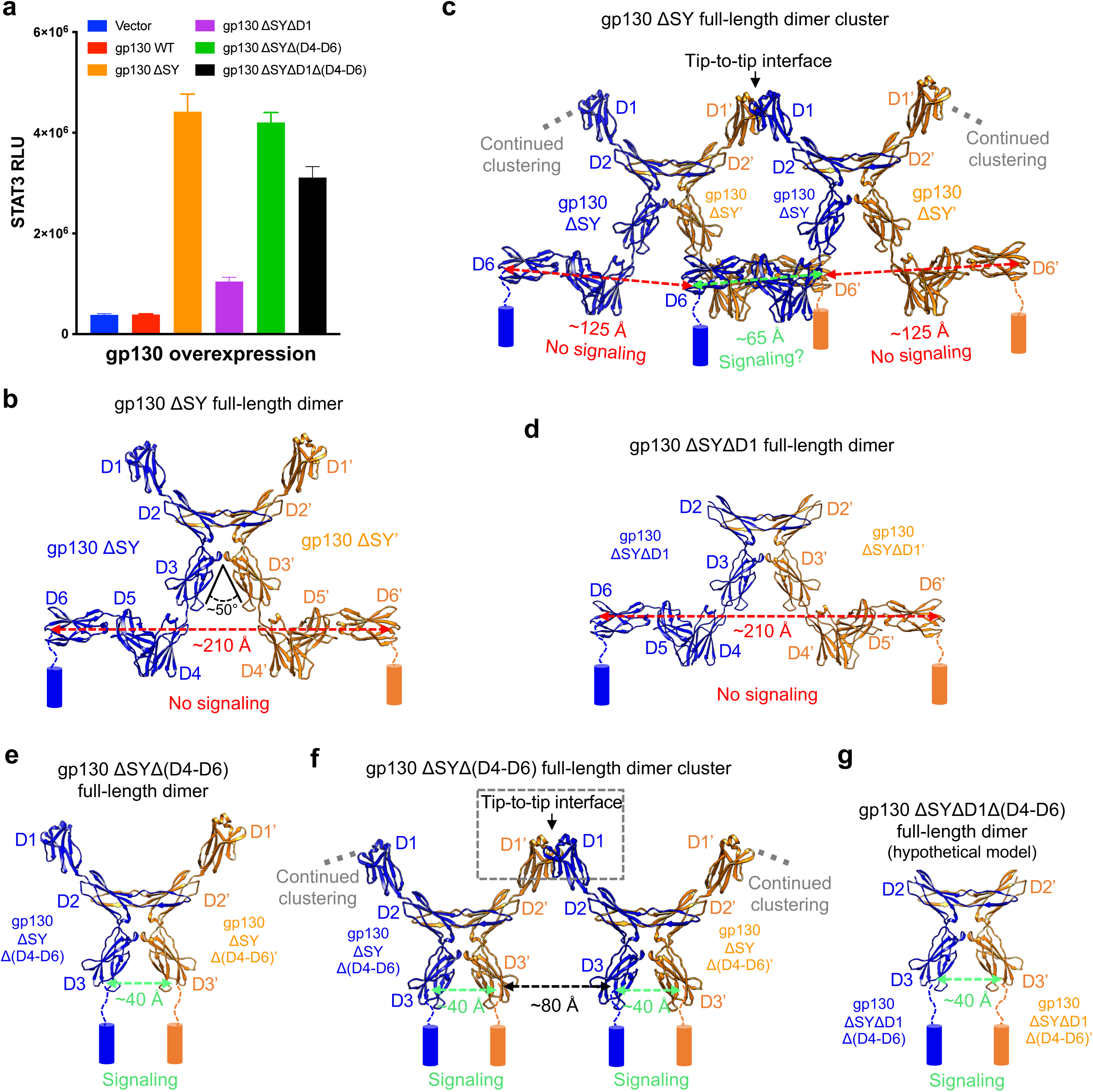
Clustering of the full-length gp130 S187-Y190 deletion mutant (gp130 ΔSY full-length) dimer via tip-to-tip interactions of D1 is critical for its constitutive activation. **a** STAT3-luciferase reporter assay was performed in 293T cells after transiently overexpressing gp130 WT or mutants as indicated. The data shown are representative of 3 independent experiments. RLU: relative luminescence units. **b** Side view of cryo-EM structure of full-length gp130 ΔSY dimer. The two gp130 ΔSY molecules are shown in blue and orange, respectively. The approximate distance between the two juxtamembrane D6 domains is estimated. Cylinders represent TM domains. **c** Side view of cryo-EM structure of full-length gp130 ΔSY dimer clusters shows clustering of the dimer mediated by tip-to-tip interactions of D1. **d** Side view of cryo-EM structure of full-length gp130 ΔSYΔD1 dimer. **e** Side view of cryo-EM structure of full-length gp130 ΔSYΔ(D4-D6) dimer. **f** Side view of cryo-EM structure of full-length gp130 ΔSYΔ(D4-D6) dimer clusters demonstrates clustering of the dimer via tip-to-tip interactions of D1. **g** Hypothetical model of full-length gp130 ΔSYΔD1Δ(D4-D6) dimer.

We hypothesized that the absence of the transmembrane (TM) helix and cytoplasmic region in the gp130 ΔSY ECD protein might result in the loss of key structural information necessary for elucidating the molecular mechanism behind mutant activation. To examine this, we purified the full-length gp130 ΔSY mutant in detergent (Supplementary Fig. 6a-b). Native PAGE gel analysis revealed that the protein associated with the main peak consisted of both monomeric and dimeric forms. Additionally, a distinct smear observed above the dimer band likely indicates the presence of protein aggregates or clustered dimers (Supplementary Fig. 6c). The sample was further examined using cryo-EM in the presence of B-S12 Fab, revealing a 3.29 Å resolution “X”-shaped dimer structure identical to that observed for the gp130 ΔSY ECD protein (Fig. 4b, Supplementary Fig. 7a-e). Notably, the TM and cytoplasmic domains were both not resolved, likely due to high flexibility of gp130 beyond the ECD region^9,10^.

Upon closer examination of the 2D class averages of particles in the top-down/bottom-up views, we noticed additional fuzzy density on both sides of the dimer, which likely signifies ongoing clustering of the dimer (Supplementary Fig. 7f). To further investigate, the particles were re-extracted using a 2.1x larger box size, revealing clear evidence of dimer clustering in the resulting new 2D class averages (Supplementary Fig. 7g). 3D reconstruction of the re-extracted particles, together with particles from a tilted dataset, produced a cryo-EM map depicting a dimer-of-dimer structure of full-length gp130 ΔSY (Supplementary Fig. 8). Interestingly, the two dimers are connected through tip-to-tip interactions, where the D1 domain of one dimer contacts D1 of another dimer (Fig. 4c). The additional densities flanking the two dimers suggest ongoing clustering (Supplementary Fig. 8d), highlighting the potential of gp130 ΔSY dimer to assemble into a one-dimensional array structure on the membrane. In this cluster, the juxtamembrane D6 domains of two interacting gp130 molecules from neighboring dimers are ∼65 Å apart (Fig. 4c), a distance substantially shorter than that observed in isolated dimers (∼210 Å, Fig. 4b) but still greater than the distance between D6 domains of two WT gp130 molecules in the IL-6 receptor complex (∼26 Å, Fig. 3g). In cells, the membrane environment might reduce this distance by leveraging potential TM or cytoplasmic interactions and the inherent flexibility of gp130 at its domain-domain hinge regions. Alternatively, two interacting gp130 molecules from adjacent dimers may possibly initiate weak signaling, which could be amplified through further clustering of the dimers^23^.

To validate the pivotal role of D1 in gp130 ΔSY dimer clustering, we purified a D1 truncated variant of the protein (Supplementary Fig. 9a) and analyzed its structure using B-S12 and 16683 Fabs as fiducial markers (Supplementary Fig. 10a-e). As expected, the removal of D1 does not affect the dimerization of this variant but eliminates its ability to cluster (Fig. 4d, Supplementary Fig. 10f), consistent with the finding that D1 is essential for constitutive activation but dispensable for dimerization of gp130 ΔYY^13,16^. In contrast, full-length gp130 ΔSY exhibits dimer clustering in the presence of B-12 and 16683 Fabs (Supplementary Fig. 10g).

The FNIII domains (D4-D6) of gp130 are not involved in binding to IL-6 and IL-6Rα^11^; however, they are essential for bringing the two juxtamembrane D6 domains into close proximity (Fig. 3g), making them critical for IL-6-induced gp130 activation^24,25^. Interestingly, truncation of D4-D6 in gp130 ΔSY showed no significant effect on its ligand-independent activity (Fig. 4a). A D4-D6 deleted variant of gp130 ΔSY was purified (Supplementary Fig. 9b) and characterized by cryo-EM, using B-S12 as a fiducial marker (Supplementary Fig. 11a-e). This analysis revealed a dimeric structure of the variant, exhibiting a shortened “X” shape (Fig. 4e). Similar to the full-length gp130 ΔSY sample, gp130 ΔSYΔ(D4-D6) particles re-extracted with a 2.25x larger box size resulted in 2D class averages and a 3D map that clearly demonstrated dimer clustering via “tip-to-tip” interactions of D1 domains from neighboring dimers (Supplementary Fig. 11f, 12 and Fig. 4f). The presence of additional weak densities on both sides of the two dimers indicates continued clustering of the dimers to form one-dimensional array structure (Supplementary Fig. 12d). Upon truncation of D4-D6, D3 becomes the new juxtamembrane domain of the variant, positioned ∼40 Å from the opposing D3 in each dimer (Fig. 4e, f), a distance that might be close enough to trigger downstream signaling (Fig. 4a). In comparison, while the deletion of domains D4-D6 from WT gp130 allows it to still form a complex with IL-6 and IL-6Rα^11^, it fails to initiate signaling (Supplementary Fig. 13a), likely because the ∼90 Å separation between the two new juxtamembrane D3 domains in the IL-6/IL-6Rα/gp130 Δ(D4-D6) complex is too large to support intracellular JAK dimerization and activation (Supplementary Fig. 13b).

Notably, further truncation of D1 from the gp130 ΔSYΔ(D4-D6) variant, which should eliminate clustering of this variant dimer, results in only a slight reduction in its constitutive activity (Fig. 4a). This result confirms that signaling can be initiated directly by the two truncated gp130 molecules within each dimer (Fig. 4g), while the clustering of dimers may further enhance the signaling. From another perspective, deleting D4-D6 from the gp130 ΔSYΔD1 variant reduces the juxtamembrane domain distance from ∼210 Å to ∼40 Å (Fig. 4d, g), largely restoring the lost activity of this variant (Fig. 4a). Collectively, these data underscore the crucial role of juxtamembrane domain proximity in gp130 activation.

In the cryo-EM maps of the two dimer clusters we obtained (Supplementary Fig. 8d, 12d), the resolutions are insufficient to resolve the detailed “tip-to-tip” interactions of D1. Nevertheless, the glycan densities observed at two glycosylation sites, N43 and N83, enable accurate docking of published D1 structure into the map, providing a general overview of how the two D1 domains interact within the dimer cluster (Supplementary Fig. 12e). The interactions are primarily mediated by residues located in the short helix and loop regions at the N-terminal tip of D1, likely through hydrophobic interactions across a relatively small interface. The fact that dimer clustering was observed in full-length gp130 ΔSY and full-length gp130 ΔSY with D4-D6 truncation, but not in gp130 ΔSY ECD, suggests that the “tip-to-tip” interaction is weak and relies on the presence of TM and cytoplasmic domains. Together with the N-terminal “tip-to-tip” interaction, these domains may stabilize the dimer clustering through interactions at the C-terminal end. Notably, a similar “tip-to-tip” geometry has been observed in the IL-17 ligand-receptor complexes^26^.

### 3D domain swapping mediates dimerization of three additional IHCAs-associated gp130 mutants with mutation in D2

As previously noted, the disruption of EF loop positioning within D2 by engineered or IHCAs-associated mutations can induce domain swapping-mediated dimerization of gp130. To further explore whether this mechanism applies to other gp130 mutants identified in IHCAs, we purified three additional representative mutants in detergent: the K173-D177 deletion mutant (ΔKD, Supplementary Fig. 9c), the E195-V196 deletion mutant (ΔEV, Supplementary Fig. 9d), and the P216H mutant (Supplementary Fig. 9e), all of which exhibit constitutive activity in cells^1^. The K173-D177 and E195-V196 regions flank the EF loop and are situated in the C’E loop and β_F_ sheet of D2, respectively (Fig. 2a). The P216 residue, on the other hand, resides in the D2D3 hinge region and forms a hydrophobic core with V189, V192, and I194 in the EF loop (Fig. 2a). Theoretically, all three mutants should have disrupted EF loop positioning, leading to 3D domain swapping and dimerization.

Cryo-EM analysis was performed on these gp130 mutants using B-S12 Fab alone or in combination with 16683 Fab to facilitate structural determination (Supplementary Fig. 14-16). As expected, all three mutants exhibit “X”-shaped dimer configurations, with the TM and cytoplasmic domains remaining unresolved due to flexibility (Fig. 5a-i). In these dimers, the β_C’_-C’E loop-β_E_ module in D2 of each monomer adopts an extended, unfolded conformation, interacting with the opposing D2 instead of folding back against its own D2. The structural differences among the dimers are notable. The gp130 ΔEV dimer exhibits an ∼50° angle between its two D3 domains (Fig. 5d), similar to the angle observed in the gp130 ΔSY dimer (Fig. 4b). In contrast, the ΔKD and P216H dimers exhibit significantly larger angles of approximately 100° and 90° (Fig. 5a, g), respectively. The spatial orientation of the two gp130 molecules within each dimer differs. In the gp130 ΔEV dimer, both molecules align nearly within the same plane, creating a relatively flat configuration (Fig. 5f), which is comparable to the ΔSY and D215G dimers (Fig. 1c, 3c). On the other hand, the gp130 molecules in the ΔKD and P216H dimers are rotated in opposite directions, diverging from the plane of the dimer interface (Fig. 5c, 5i), resembling the arrangement observed in the V252G dimer (Fig. 3f). These variations in angle and planar configuration lead to distinct separations between the juxtamembrane D6 domains within the dimers: approximately 230 Å for the ΔKD dimer, 210 Å for the ΔEV dimer, and 245 Å for the P216H dimer (Fig. 5a, d, g). These distances are evidently too large to support the dimerization and activation of JAK kinases.

**Fig. 5.**
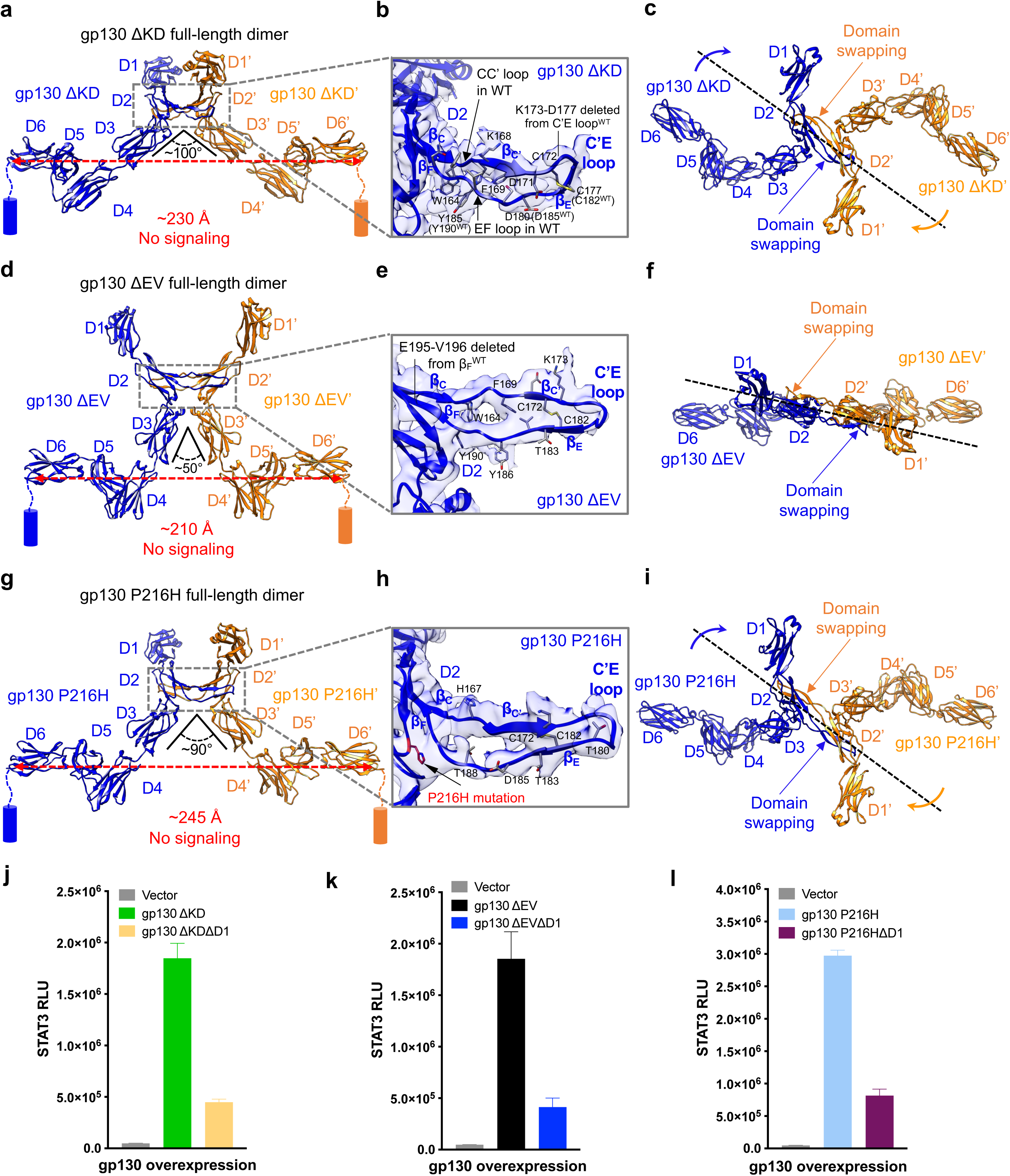
3D domain swapping mediates dimerization of three additional IHCAs-associated gp130 mutants with deletion or point mutation in D2. **a** Side view of cryo-EM structure of the gp130 K173-D177 deletion mutant (gp130 ΔKD) dimer. The two gp130 ΔKD molecules are colored in blue and orange, respectively. Approximate distance between the two D6 domains is estimated. Cylinders represent TM domains. **b** Zoomed-in view of atomic model in transparent density of D2 of the blue gp130 ΔKD molecule shows an extended conformation of the β_C’_-C’E-β_E_ module. A few residues in the module are shown in stick representation. **b** is in the same orientation as **a**. **c** Top-down view of cryo-EM structure of the gp130 ΔKD dimer. Dash line indicates the plane of the dimer interface. Arrows indicate the tilting of the two gp130 molecules away from the plane of the dimer interface. **d** Side view of cryo-EM structure of the gp130 E195-V196 deletion mutant (gp130 ΔEV) dimer. **e** Zoomed-in view of atomic model in transparent density of D2 of the blue gp130 ΔEV molecule shows an extended conformation of the β_C’_-C’E-β_E_ module. **e** is in the same orientation as **d**. **f** Top-down view of cryo-EM structure of the gp130 ΔEV dimer. Dash line indicates the plane of the dimer interface. **g** Side view of cryo-EM structure of the gp130 P216H mutant dimer shows 3D domain swapping of the β_C’_-C’E-β_E_ module in D2. **h** Zoomed-in view of atomic model in transparent density of D2 of the blue gp130 P216H molecule shows an extended conformation of the β_C’_-C’E-β_E_ module. **h** is in the same orientation as **g**. **i** Top-down view of cryo-EM structure of the gp130 P216H dimer. Dash line indicates the plane of the dimer interface. Arrows indicate the tilting of the two gp130 molecules away from the plane of the dimer interface. **j** STAT3-luciferase reporter assay was performed in 293T cells after transiently overexpressing gp130 ΔKD or gp130 ΔKDΔD1. The data shown are representative of 3 independent experiments. RLU: relative luminescence units. **k** STAT3-luciferase reporter assay was performed in 293T cells after transiently overexpressing gp130 ΔEV or gp130 ΔEVΔD1. The data shown are representative of 3 independent experiments. **l** STAT3-luciferase reporter assay was performed in 293T cells after transiently overexpressing gp130 P216H or gp130 P216HΔD1. The data shown are representative of 3 independent experiments.

We hypothesized that the constitutive activities of these gp130 mutants are also driven by D1-mediated dimer clustering, similar to gp130 ΔSY. Consistently, the deletion of D1 from these mutants significantly reduced ligand-independent activation across all three mutants (Fig. 5j-l). Furthermore, clear evidence of dimer clustering was observed in the 2D averages of the ΔEV sample (Supplementary Fig. 15f), though not for the ΔKD and P216H samples. Notably, dimer clustering was apparent only in 2D averages of ΔSY and ΔEV particles with top-down/bottom-up view (Supplementary Fig. 7g, Supplementary Fig. 15f), likely because dimer clusters with side orientations were less stable and fell apart at the air-water interface on the EM grid. The ΔKD and P216H particles exhibited a clear orientation preference, with nearly no top-down or bottom-up view (Supplementary Fig. 14a, Supplementary Fig. 16a), which may explain why there is no dimer clustering observed in these two samples.

### 3D domain swapping drives dimerization and activation of IHCAs-associated gp130 A418-F421 deletion mutant

In addition to the IHCAs-associated mutations observed at the gp130 D2 hot spot, a rare in-frame deletion involving four residues (A418-F421) was identified at the D4D5 junction. This deletion has been shown to promote homodimerization and trigger constitutive activation of gp130^1^. However, the mechanism underlying this activation remains poorly understood.

We purified this mutant (gp130 ΔAF) in detergent (Supplementary Fig. 9f) and determined its cryo-EM structure using the 16683 Fab as a fiducial marker (Supplementary Fig. 17a-e). As anticipated, this mutant exists as a homodimer with 2-fold symmetry. The 16683 Fab binds to the D5 domain of gp130 ΔAF on the side opposite to the dimer interface and is not involved in the dimerization (Supplementary Fig. 17f). Notably, rather than adopting the “X”-shaped, “back-to-back” dimer conformation with separated juxtamembrane domains typically observed in gp130 variants with mutations in D2, the gp130 ΔAF dimer exhibits a distinct “knee-to-knee” arrangement. In this configuration, the membrane-proximal domains (D5D6) of the two monomers align parallel to one another, creating a structure that resembles a “Y” shape (Fig. 6a-b). This dimerization is also mediated by a 3D domain-swapping mechanism. However, unlike other gp130 variants where only a small module within the D2 domain is swapped, the gp130 ΔAF dimer involves the exchange of the entire D5D6 domains between the two monomers (Fig. 6a-b). In WT gp130, the D5 domain folds back against D4, forming an acute bend (∼80°) at the D4D5 hinge region. However, the deletion of four residues (A418-F421) in gp130 ΔAF shortens the hinge, preventing D5 from bending. Consequently, D5 in the ΔAF variant rotates ∼100° and adopts an extended conformation (Fig. 6c-e). Instead of interacting with D4 of the same chain through hydrophobic interactions, the D5 domain in each gp130 ΔAF monomer packs against the D4 domain of the opposing chain, preserving the critical inter-domain interactions seen in WT gp130 (Supplementary Fig. 18)^27^.

**Fig. 6.**
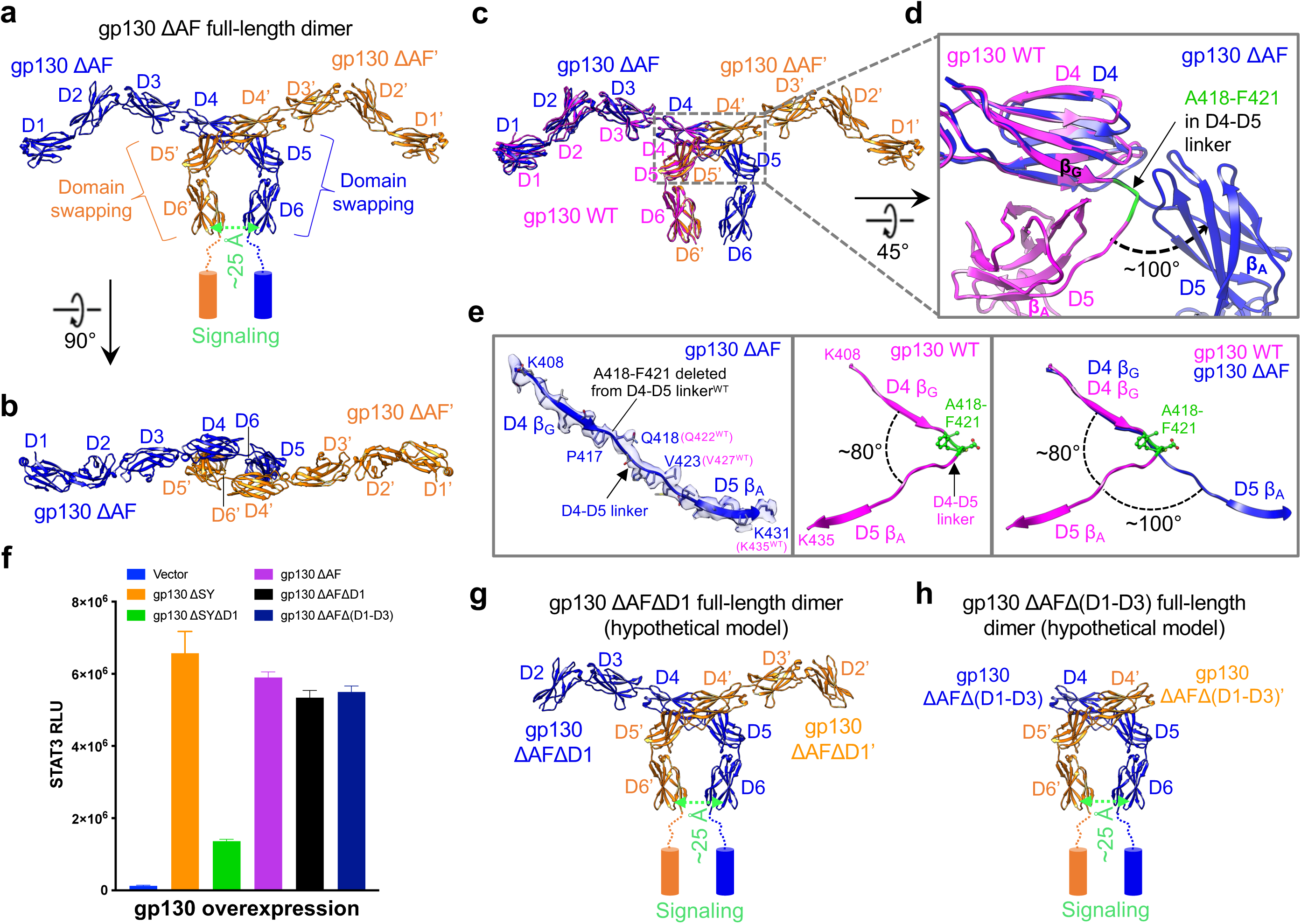
3D domain swapping mediates dimerization of IHCAs-associated gp130 A418-F421 deletion mutant (gp130 ΔAF). **a** Side view of full-length gp130 ΔAF dimer structure shows swapping of D5D6 domains between the two gp130 ΔAF molecules colored in blue and orange, respectively. The distance between the two D6 domains is ∼25 Å. Cylinders represent TM domains. **b** Top-down view of gp130 ΔSY ECD dimer structure. **c** Overlay of gp130 ΔAF dimer structure (blue and orange) with gp130 WT structure (magenta; PDB: 86D2). **d** Zoomed-in view of the overlaid structures in **c** shows a ∼100° rotation of gp130 ΔAF D5D6 relative to gp130 WT D5D6 upon deletion of A418-F421 located in the D4D5 linker region. **d** is rotated 45° around x-axis relative to **c**. gp130 ΔAF’ molecule colored in orange in **c** is not shown in **d** for better visibility. **e** Left: zoomed-in view of atomic model in transparent density of K408-K431 of the blue gp130 ΔAF molecule shows an extended conformation of D4 β_G_ and D5 β_A_ in the mutant. Residues in this fragment are shown in stick representation; middle: the K408-K435 fragment in gp130 WT shows an acutely bent (∼80°) conformation of D4 β_G_ and D5 β_A_; right: overlay of the left and middle panels shows a ∼100° rotation of D5 β_A_ in gp130 ΔAF compared to gp130 WT. **e** is in the same orientation as **a**. **f** STAT3-luciferase reporter assay was performed in 293T cells after transiently overexpressing gp130 WT or various gp130 mutants as indicated. The data shown are representative of 3 independent experiments. RLU: relative luminescence units. **g** Hypothetical model of full-length gp130 ΔAFΔD1 dimer. **h** Hypothetical model of full-length gp130 ΔAFΔ(D1-D3) dimer.

Remarkably, the two juxtamembrane D6 domains in the gp130 ΔAF dimer are positioned approximately 25 Å apart (Fig. 6a), a distance comparable to that observed in the IL-6 receptor complex (∼26 Å, Fig. 3g). This indicates that the gp130 ΔAF dimer may already be active without requiring the formation of dimer clusters through “tip-to-tip” interactions of D1, as seen in other IHCAs-associated gp130 variants with mutations in D2. In fact, truncating D1 had minimal impact on the activity of gp130 ΔAF, nor did the deletion of the entire D1-D3 domains (Fig. 6f), suggesting that these deletions would not alter the dimerization status or the proximity of the juxtamembrane D6 domains (Fig. 6g, h). Although we did not directly observe evidence of gp130 ΔAF dimer clustering on the EM grid (Supplementary Fig. 17a), we cannot rule out the possibility that dimers may still cluster via interaction mediated by D1 of neighboring dimers on the membrane. Nevertheless, such clustering is not essential for the constitutive activation of gp130 ΔAF.

## Discussion

The constitutive activation of gp130 in IHCAs is primarily driven by gain-of-function mutations that compromise the structural integrity or positioning of the EF loop within domain D2, which appears to be crucial for maintaining the folded conformation of the β_C’_-C’E-β_E_ module in D2. The most common gp130 mutation in IHCAs, the S187-Y190 deletion (gp130 ΔSY), shortens the EF loop, causing the β_C’_-C’E-β_E_ module to unfold. Rather than packing against its own D2 domain, this module interacts with D2 of the opposing gp130 ΔSY molecule, facilitating dimerization of the mutant. Other IHCAs-associated mutants, such as ΔKD (K173-D177 deletion), ΔEV (E195-V196 deletion), and P216H, share a similar hallmark of EF loop alteration or destabilization. These mutations either shorten the regions flanking the loop (ΔKD and ΔEV) or disrupt hydrophobic interactions critical for proper loop positioning (P216H). Consequently, these mutants also exhibit domain-swapping-mediated dimerization. In addition, engineered mutations like D215G and V252G, which impair hydrogen bonding and hydrophobic interactions responsible for stabilizing the EF loop and the folded conformation of the β_C’_-C’E-β_E_ module within D2, recapitulate the domain-swapping mechanisms observed in IHCAs-associated mutants. Notably, a D215 deletion variant has been identified in IHCAs^1^, highlighting the critical role of the D215 residue in preserving the native conformation of gp130.

All these mutants adopt an “X”-shaped dimer configuration, where the two monomers are arranged in a “back-to-back” orientation. In this arrangement, the D2D3 elbow regions of each gp130 molecule lean against one another, held together by interactions between the swapped β_C’_-C’E-β_E_module and D2 of the opposing gp130 molecule. Despite the shared structural feature, the β_C’_-C’E-β_E_ modules exhibit diverse conformations influenced by specific structural alterations in different mutants, which lead to various dimer configurations (Supplementary Fig. 19). The ΔSY, ΔEV, and D215G mutants create dimers with a compact inter-domain angle of 30-50° between their D3 domains. In these dimers, the gp130 molecules are arranged nearly in a flat, planar structure, with the juxtamembrane D6 domains separated by a distance of 200-210 Å. In contrast, the ΔKD, P216H, and V252G mutants generate dimers with much wider D3-D3 angles ranging from 90-100°. These dimers exhibit a distinct structural configuration where the gp130 molecules tilt away from the plane of the dimer interface in opposite directions, breaking the planar alignment. This divergence results in an expanded distance between the D6 domains, which increases to 230-250 Å.

Based on AlphaFold prediction and prior research^28,29^, gp130 structure is characterized by short linker regions, about 10 residues in length, between D6 and the TM domain, as well as between TM and the intracellular Box1/Box2 JAK-binding motif. These short linkers impose structural constraints that necessitate the juxtamembrane D6 domains to be in close proximity to enable JAK dimerization and activation, which is contingent upon an approximately 50 Å distance between the centers of the JAK binding sites on the two receptors (Fig. 7a)^19,20^. Consistent with this requirement, the IL-6/IL-6Rα/gp130 signaling complex adopts an “O”-shaped configuration, with the juxtamembrane domains positioned at a close distance (∼26 Å). In contrast, the “X”-shaped open dimer structures observed in these gp130 variants, characterized by juxtamembrane domain separations exceeding 200 Å, are highly unexpected, as such separations would physically preclude JAK kinase dimerization and activation (Fig. 7b)^19–22^. However, our cryo-EM data on full-length gp130 ΔSY suggest that the clustering of ΔSY dimers may compensate for such structural limitation of the isolated dimer, enabling signaling between two gp130 molecules from neighboring dimers connected via “tip-to-tip” interactions of D1. This signaling is probably weak due to the ∼65 Å distance at the juxtamembrane D6 domain (Fig. 7c), which is larger than the distance observed in the IL-6 receptor complex. However, it is possible that further clustering of receptor dimers may amplify the signaling (Fig. 7b)^30–33^. Importantly, gp130 ΔSY dimer clustering appears to be context-dependent, requiring the presence of transmembrane and cytoplasmic domains. The observation of dimer clustering in full-length gp130 ΔSY and full-length gp130 ΔSY with D4-D6 truncation, but not in gp130 ΔSY ECD, highlights that the “tip-to-tip” interaction is probably inherently weak and relies on the TM and cytoplasmic domains for stabilization. The membrane anchoring and potential C-terminal interactions mediated by these domains, along with the N-terminal “tip-to-tip” interaction, likely stabilize dimer clustering in a one-dimensional array configuration. Notably, a similar “tip-to-tip” receptor interaction has been observed in IL-17 ligand-receptor complexes, where the interaction itself is weak and depends on the initial binding of the ligand^26^.

**Fig. 7.**
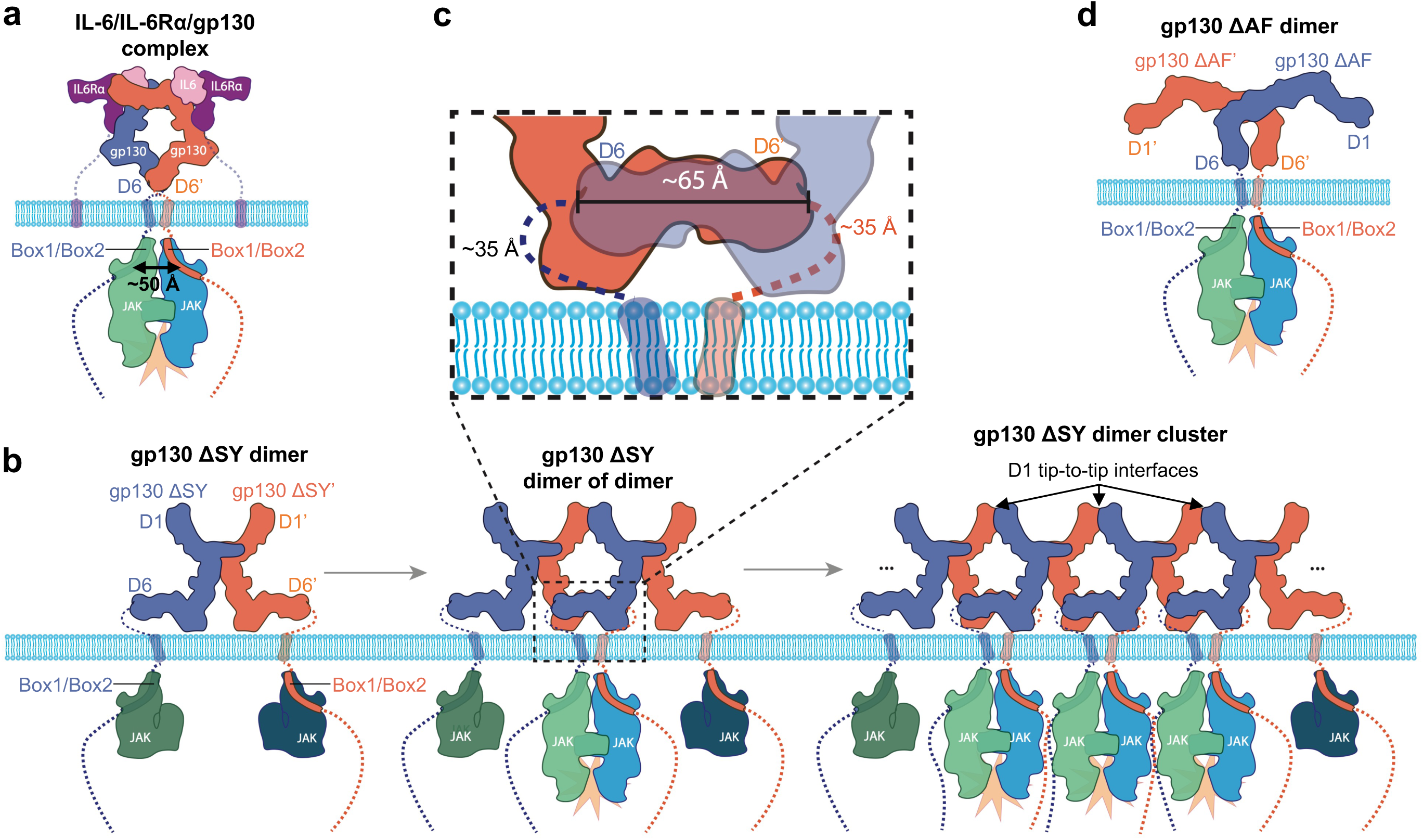
Juxtamembrane domain proximity is crucial for both ligand-dependent and mutation-induced activation of gp130. **a** Signaling model of the IL-6 receptor complex. Cylinders represent TM domains. Flexible regions of gp130 and IL-6Rα are shown in dash lines. The distance between the centers of the JAK binding sites (Box1/Box2 motifs) on the two gp130 receptors is estimated. **b** Signaling model of the gp130 ΔSY mutant. **c** Zoomed-in view of boxed region from **b**. The distance between the two juxtamembrane D6 domains from two adjacent dimers, as well as the length of the linker region between D6 and TM, are estimated. **d** Signaling model of the gp130 ΔAF mutant.

Overexpression of WT gp130 has been shown to inhibit gp130 ΔSY activity in a dose-dependent manner in cells^4^, likely by blocking ΔSY dimer clustering through “tip-to-tip” interactions of D1 domains at both sides of the dimer (Supplementary Fig. 20a). Interestingly, the soluble ECD protein of WT gp130 does not affect the constitutive activity of gp130 ΔYY, another common variant associated with IHCAs^16^. This is probably because the TM and cytoplasmic domains, which appear to be essential for stabilizing the weak D1-D1 interaction-mediated clustering, are absent in the ECD protein. Moreover, previous studies suggest that WT gp130 can exist as a homodimer in the absence of ligand on the cell membrane^34,35^, potentially driven by “tip-to-tip” interactions of the D1 domain (Supplementary Fig. 20b), similar to the D1-D1 interactions seen in gp130 ΔSY dimer cluster. However, due to the lack of structural stabilization provided by active dimer clustering, the WT gp130 dimer is probably only transiently formed on the membrane^34^, which may impede its ability to initiate signaling. Furthermore, the relatively large distance between the juxtamembrane D6 domains may be another factor contributing to the inactivity of the pre-formed WT gp130 dimer in the absence of a ligand.

Therapeutically, the ability of WT gp130 to inhibit the constitutive activity of certain mutant, such as gp130 ΔSY, offers a promising avenue for intervention. In addition, the identification of the D1 domain as a critical mediator of dimer clustering provides a potential target for therapeutic modulation of gp130 variants requiring dimer clustering for constitutive activation. Anti-gp130 antibodies disrupting the dimer clustering but not impairing cytokine-receptor complex assembly may selectively inhibit these variants without affecting normal gp130 signaling. However, the findings that one of the variants, gp130 ΔYY, is mainly retained intracellularly and transduces signal from intracellular compartments^8,36^, suggest that targeting the mutant receptor externally using antibodies may not efficiently inhibit all aberrant gp130 activation. Instead, membrane permeable blockers of dimer clustering, such as small molecules, peptides, helicons, etc., may provide a more effective therapeutic strategy. Furthermore, therapies targeting the intracellular signaling cascades, such as the JAK1/JAK2 inhibitor Ruxolitinib, have been proposed for the treatment of IHCAs expressing constitutively active gp130 mutants and certain subsets of HCCs with similar mutations^1^.

While domain swapping is a common feature, IHCAs-associated gp130 mutants display structural diversity in their dimer configurations. For instance, in contrast to the “X”-shaped “back-to-back” dimer conformations observed in variants with mutations in D2, where the β_C’_-C’E-β_E_ module in D2 undergoes swapping, a unique A418-F421 deletion (gp130 ΔAF) at the D4D5 junction adopts a distinct “Y”-shaped “knee-to-knee” dimer arrangement. This configuration involves the exchange of entire D5D6 domains between monomers, aligning the D5D6 domains in parallel and bringing the juxtamembrane domains into close proximity (∼25 Å). This proximity enables constitutive activation without requiring dimer clustering (Fig. 7d). These structural variations highlight the heterogeneity of gp130 activation mechanisms and suggest that different mutations may utilize distinct strategies to achieve constitutive signaling.

In conclusion, our study provides a comprehensive analysis of gp130 activation mechanisms, focusing on the structural and functional consequences of somatic gp130 mutations associated with IHCAs. By elucidating the molecular basis for the dimerization and activation of gp130 mutants, our findings bridge critical gaps in understanding the pathogenesis of IHCAs and offer avenues for targeted therapeutic strategies.

## Methods

### Expression and purification of soluble extracellular domain of gp130

In this study, protein residue numbering includes the signal peptides. The soluble extracellular domain (ECD, residues E23-T619) of gp130 wild-type (WT), S187-Y190 deletion mutant (ΔSY), D215G mutant, and V252G mutant were expressed as C-terminal myc-myc-His (mmH)-tagged secreted proteins in HEK-293 (Expi293F) cells following transfection with Lipofectamine 2000 (Thermo Fisher, #11668027). To purify each protein, the filtered cell culture supernatant was buffer exchanged against DPBS via dialysis and loaded onto a pre-equilibrated Talon column (Clontech, #635682). The column was washed first with DPBS containing 500 mM NaCl, followed by a second wash with DPBS supplemented with 500 mM NaCl and 5 mM Imidazole. Proteins were eluted using DPBS containing 500 mM NaCl and 200 mM Imidazole, and the eluates were further purified via size exclusion chromatography (SEC) using a Superdex 200 Increase 10/300 GL column (Cytiva).

### Expression and purification of full-length gp130

The full-length gp130 mutants analyzed in this study include the gp130 S187-Y190 deletion mutant (ΔSY), gp130 ΔSY with an additional D1 deletion (gp130 ΔSYΔD1), gp130 ΔSY with additional D4-D6 deletion (gp130 ΔSYΔ(D4-D6)), gp130 K173-D177 deletion mutant (gp130 ΔKD), gp130 E195-V196 deletion mutant (gp130 ΔEV), and gp130 P216H mutant. All of these were C-terminal Flag-tagged proteins purified in detergent. To express these proteins, the genes of interest were synthesized and cloned into the pEZT-BacMam vector by GenScript. The plasmid was transformed into MAX Efficiency DH10Bac Competent Cells (Life Technologies, 10361012) to generate recombinant Bacmid DNA containing the respective genes, which were subsequently used to transfect ExpiSF9 Insect cells (Life Technologies, A35243) cultured in ExpiSf CD Medium (Life Technologies, A3767803) to produce Baculovirus. After 3 days of culture, the cells were pelleted by centrifugation, and the virus-containing supernatant was collected and subjected to ultracentrifugation. The viral pellet was then resuspended in Freestyle 293 Expression Medium (Gibco, 12338-018) containing 2% Fetal Bovine Serum (Life Technologies, A3840101) and filtered for transduction into Freestyle HEK 293-F cells (Life Technologies, R79007) cultured in Freestyle 293 Expression Medium. Sodium Butyrate (10 mM, Thermo Scientific, 263191000) was added 24 hours post-transduction to enhance transduction efficiency. The culture was grown for an additional 24 hours, harvested by centrifugation, and the cell pellet was washed with pre-chilled DPBS before being frozen at -80°C.

For protein purification, each cell pellet was resuspended in solubilization buffer (20 mM HEPES pH 7.4, 150 mM NaCl, 1% Lauryl Maltose Neopentyl Glycol (LMNG, Anatrace, #NG310), cOmplete EDTA-free protease inhibitors, and Benzonase Nuclease) and rotated at 4°C for 1 hour. After centrifugation, the supernatant was incubated with anti-Flag M2 affinity resin (Sigma Aldrich, #A2220) pre-equilibrated with SEC buffer (20 mM HEPES pH 7.4, 150 mM NaCl, 0.01% LMNG, 0.001% CHS) at 4°C for 30 minutes. The resin was then washed with SEC buffer, and the protein was eluted using SEC buffer supplemented with 0.15 mg/ml of 3X Flag peptide. The eluates were further purified by size exclusion chromatography (SEC). Specifically, the full-length gp130 ΔSYΔ(D4-D6) mutant was purified using a Superdex 200 Increase 10/300 GL column, while the other mutants were purified using a Superose 6 Increase 10/300 GL column. Peak fractions were collected and concentrated using a 100-kDa molecular weight cutoff (MWCO) centrifugal concentrator.

### Fab fragment preparation

The Fab fragment of the anti-gp130 antibody B-S12 (Abcam, #ab27359) was prepared using the Pierce™ Mouse IgG1 Fab and F(ab’)2 Preparation Kit (Thermo Fisher, #44980) following the manufacturer’s instructions. The anti-gp130 antibody H4xH16683P2 was digested into F(ab’)2 and Fc fragments using the Fabricator enzyme (Genovis). The F(ab’)2 fragment was further reduced into F(ab)’ using 2-mercaptoethylamine (2-MEA, Thermo Fisher), followed by the removal of Fc fragments using the CaptureSelect IgG-Fc (ms) affinity resin (Thermo Fisher). Each antibody’s Fab fragment was subsequently purified using a Superdex 200 Increase 10/300 GL gel filtration column equilibrated with 50 mM Tris (pH 7.5) and 150 mM NaCl. The purified Fab fragments were concentrated using a 10 kDa MWCO centrifugal concentrator.

### gp130-Fab complex preparation

The soluble ECD protein of gp130 WT or mutant was mixed with B-S12 Fab alone or with both B-S12 Fab and H4xH16683P2 Fab in an equimolar ratio, followed by incubation at 4°C for 30 minutes. Each mixture was then concentrated to approximately 3 mg/ml using a 30 kDa MWCO centrifugal concentrator. Similarly, full-length gp130 mutant proteins (∼0.8 mg/ml), solubilized in detergent, were incubated with B-S12 Fab, H4xH16683P2 Fab, or both in an equimolar ratio at 4°C for 30 minutes.

### Cryo-EM grid preparation and data collection

Each gp130 ECD/Fab complex was mixed with approximately 0.15% Amphipol PMAL-C8 (Anatrace) immediately prior to applying 3.5 µL of the mixture onto an UltrAufoil R0.6/1, 300-mesh grid (Quantifoil). For each full-length gp130 mutant/Fab complex, 3.5 µL was directly applied to the grid without Amphipol. The grids were blotted for 4.5 seconds at a force of 0 and plunge-frozen into liquid ethane using a Vitrobot Mark IV (ThermoFisher) operated at 4°C and 100% humidity. The frozen grids were loaded into a Titan Krios G3i microscope (ThermoFisher) equipped with a K3 BioQuantum electron detector and energy filter (Gatan) for data acquisition at a nominal magnification of 105,000x. Automated data collection was performed using EPU v2.12 (ThermoFisher) with a defocus range of -1.2 to -2.6 µm and an energy filter slit width of 20 eV. Each movie was dose-fractionated into 46 frames over a 2-second exposure with a total dose of 40 electrons/Å^2^.

### Cryo-EM data processing

Cryo-EM data were processed using cryoSPARC v2^37^. Movies were motion corrected by Patch motion correction and CTF parameters were estimated by Patch CTF estimation. Particles were initially picked using Blob picker from ∼500 micrographs to generate 2D class averages, which were used as templates for the subsequent template picking. Cryo-EM data processing statistics were summarized in Supplementary Table 1.

For the gp130 ΔSY ECD/B-S12 Fab/16683 Fab complex (Supplementary Fig. 2a), 7,330,843 particles were picked from 11,094 micrographs by template picking. After 2D classification, 715,803 particles remained and were subjected to Ab initio Reconstruction and Heterogeneous Refinement, isolating a best class of 332,112 particles. A subsequent 2D classification further refined this to 291,116 particles. Homogeneous Refinement and masked Local Refinement with enforced C2 symmetry produced a map at a nominal resolution of 3.24 Å.

For the gp130 D215G ECD/B-S12 Fab/16683 Fab complex (Supplementary Fig. 4a), 8,723,011 particles were selected from 11,086 micrographs using template picking. Following 2D classification, 648,183 particles remained and underwent Ab initio Reconstruction and Heterogeneous Refinement, yielding a best class of 279,906 particles. These particles were further refined through Homogeneous Refinement and subsequent Heterogeneous Refinement, isolating two high-quality classes comprising 220,794 particles in total. Final Homogeneous Refinement and masked Local Refinement with enforced C2 symmetry resulted in a map with a nominal resolution of 3.38 Å.

For the gp130 V252G ECD/B-S12 Fab/16683 Fab complex (Supplementary Fig. 5a), template picking identified 11,010,102 particles from 14,775 micrographs. After 2D classification, 752,581 particles remained and underwent Ab initio Reconstruction and Heterogeneous Refinement, isolating a best class containing 326,777 particles. These particles were further refined through Homogeneous Refinement, followed by Heterogeneous Refinement, resulting in two high-quality classes comprising a total of 229,453 particles. Final Homogeneous Refinement and masked Local Refinement with enforced C2 symmetry produced a map at a nominal resolution of 3.82 Å.

For the gp130 ΔSY full-length/B-S12 Fab complex (Supplementary Fig. 7a), template picking identified 7,538,941 particles, which were extracted with a box size of 326.8 Å from 10,097 micrographs. Following 2D classification, 1,082,259 particles remained and underwent Ab initio Reconstruction and Heterogeneous Refinement, isolating a best class of 388,756 particles. These particles were further refined through Homogeneous Refinement, followed by Heterogeneous Refinement, yielding two high-quality classes totaling 242,322 particles. Final Homogeneous Refinement and masked Local Refinement with enforced C2 symmetry resulted in a map at a nominal resolution of 3.29 Å. To resolve the structure of the gp130 ΔSY full-length dimer cluster, an additional set of 8,582 movies were collected with the specimen stage tilted by 40°, which were combined with the original 10,097 untilted movies for data processing (Supplementary Fig. 8a). From this combined dataset, 13,912,054 particles were picked and extracted with a box size of 688 Å. After 2D classification, 1,575,587 particles remained and underwent Ab initio Reconstruction and Heterogeneous Refinement, isolating a best class of 132,676 particles. These particles were further refined through Homogeneous Refinement and Heterogeneous Refinement, yielding one high-quality class of 69,303 particles. Final Homogeneous Refinement and masked Local Refinement resulted in a map with a nominal resolution of 6.78 Å.

For the gp130 ΔSYΔD1 full-length/B-S12 Fab/16683 Fab complex (Supplementary Fig. 10a), 13,788,347 particles were initially picked from 19,470 micrographs. After 2D classification, 648,183 particles remained and were subjected to Ab initio Reconstruction and Heterogeneous Refinement, isolating a best class of 80,299 particles. These selected particles underwent further processing through Homogeneous Refinement followed by Heterogeneous Refinement, yielding two high-quality classes comprising a total of 63,329 particles. Final Homogeneous Refinement and masked Local Refinement, with enforced C2 symmetry, resulted in a map at a nominal resolution of 3.91 Å.

For the gp130 ΔSYΔ(D4-D6) full-length/B-S12 Fab complex (Supplementary Fig. 11a), 10,688,683 particles were picked and extracted with a box size of 268.8 Å from 14,147 micrographs. Following 2D classification, 2,052,508 particles remained and were subjected to Ab initio Reconstruction and Heterogeneous Refinement, isolating a best class of 1,425,787 particles. These particles underwent further processing through Homogeneous Refinement and Heterogeneous Refinement, yielding one high-quality class containing 340,107 particles. Final Homogeneous Refinement and masked Local Refinement, with enforced C2 symmetry, produced a map with a nominal resolution of 3.04 Å. To resolve the structure of the gp130 ΔSYΔ(D4-D6) full-length dimer cluster, the same 10,688,683 particles were extracted from the 14,147 micrographs but with a larger box size of 604.8 Å (Supplementary Fig. 12a). After 2D classification, 1,023,975 particles were retained and processed through Ab initio Reconstruction and Heterogeneous Refinement, isolating a best class of 320,782 particles. These particles were further refined through Homogeneous Refinement and Heterogeneous Refinement, resulting in one high-quality class with 105,713 particles. Final Homogeneous Refinement produced a map with a nominal resolution of 6.35 Å.

For the gp130 ΔKD/B-S12 Fab complex (Supplementary Fig. 14a), 5,937,687 particles were picked from 10,906 micrographs. After 2D classification, 160,723 particles were retained and subjected to Ab initio Reconstruction, isolating a best class of 111,263 particles. These particles underwent further processing through Homogeneous Refinement and Heterogeneous Refinement, resulting in one high-quality class comprising 67,458 particles. Final Homogeneous Refinement and masked Local Refinement, with enforced C2 symmetry, produced a map with a nominal resolution of 3.36 Å.

For the gp130 ΔEV/B-S12 Fab complex (Supplementary Fig. 15a), a total of 5,329,929 particles were selected from 10,134 micrographs. After 2D classification, 734,811 particles remained and were processed using Ab initio Reconstruction and Heterogeneous Refinement, leading to the isolation of a best class containing 216,585 particles. These particles underwent further refinement through Homogeneous Refinement, followed by Heterogeneous Refinement, resulting in two high-quality classes comprising a combined total of 153,320 particles. The final steps, including Homogeneous Refinement and masked Local Refinement with C2 symmetry enforced, produced a map with a nominal resolution of 4.02 Å.

For the gp130 P216H/B-S12 Fab/16683 Fab complex (Supplementary Fig. 16a), 12,904,732 particles were picked from 19,118 micrographs. 950,503 particles were left after 2D classification, which were subjected to Ab initio Reconstruction and Heterogenous Refinement to isolate a best class with 286,403 particles. The particles were further subjected to Homogeneous Refinement, followed by Heterogenous Refinement to isolate one good class with 94,739 particles. Homogeneous Refinement and masked Local Refinement of these particles with C2 symmetry enforced yielded a map with a nominal resolution of 4.09 Å.

For the gp130 ΔAF/16683 Fab complex (Supplementary Fig. 17a), a total of 8,723,011 particles were extracted from 19,193 micrographs. After 2D classification, 980,470 particles were retained and processed through Ab initio Reconstruction and Heterogeneous Refinement, resulting in the identification of a best class containing 423,902 particles. These particles underwent additional refinement steps, including Homogeneous Refinement and another round of Heterogeneous Refinement, which yielded two high-quality classes with a combined total of 274,585 particles. Final processing steps, involving Homogeneous Refinement and masked Local Refinement with C2 symmetry applied, produced a map at a nominal resolution of 3.62 Å.

### Model building and refinement

AlphaFold^28^ predicted models of various gp130 variants, as well as the B-S12 Fab and 16683 Fab, were used as initial models for model building. A combination of picked initial models was docked into corresponding cryo-EM density map to generate a model for each complex using Fit-in-map of UCSF Chimera 1.16^38^, followed by manual modeling in Coot 0.8.9^39^. The model was subsequently real-space refined with secondary structure and non-crystallographic symmetry (NCS) restraints in Phenix 1.19^40^. The geometries of the structures were validated by using MolProbity^41^ in Phenix and the statistics were summarized in Supplementary Table 1.

### STAT3 luciferase assay

HEK-293 cells (ATCC, #CRL-1573) were plated at a density of 20,000 cells per well in 100 µl of complete DMEM medium in 96-well black, clear-bottom plates coated with Poly-D-Lysine and incubated overnight at 37°C with 5% CO . The following day, the medium was replaced with 95 µl of Opti-MEM containing 0.1% Fetal Bovine Serum (FBS), and 5 µl of a Fugene-DNA complex was added to each well. The transfection mixture included 0.5 ng of gp130 WT or variant DNA and 50 ng of pGL4.47luc2P/SIE/Hygro plasmid DNA (Promega, #E4041) in Opti-MEM, utilizing FuGENE HD Transfection Reagent (Promega, #2311). The cells were then incubated for 48 hours. On the day of the assay, the medium was replaced with 50 µl of Opti-MEM supplemented with 0.1% FBS. To minimize background luminescence, the plate was sealed with a black absorbing film. 50 µl of One-Glo EX reagent (Promega, #N1620) was added to each well. The plate was shaken for 10 minutes, and luminescence was measured using a PerkinElmer EnVision plate reader (PerkinElmer, #2105). Data analysis was conducted using GraphPad Prism software. Each experiment was repeated three times to ensure reproducibility. The data from a representative experiment were shown.

## Data availability

The cryo-EM density maps and accompanying atomic coordinates have been deposited to Electron Microscopy Data Bank (EMDB) and Protein Data Bank (PDB) with the following accession codes: gp130 ΔSY ECD/B-S12 Fab/16683 Fab complex (EMD-73011 and 9YJ5), gp130 D215G ECD/B-S12 Fab/16683 Fab complex (EMD-73012 and 9YJ6), gp130 V252G ECD/B-S12 Fab/16683 Fab complex (EMD-73013 and 9YJ7), gp130 ΔSY/B-S12 Fab complex (dimer: EMD-73014 and 9YJ8; dimer cluster: EMD-73015 and 9YJ9), gp130 ΔSYΔD1/B-S12 Fab/16683 Fab complex (EMD-73017 and 9YJA), gp130 ΔSYΔ(D4-D6)/B-S12 Fab complex (dimer: EMD-73018 and 9YJB; dimer cluster: EMD-73019 and 9YJC), gp130 ΔKD/B-S12 Fab complex (EMD-73020 and 9YJD), gp130 ΔEV/B-S12 Fab complex (EMD-73021 and 9YJE), gp130 P216H/B-S12 Fab/16683 Fab complex (EMD-73022 and 9YJF), gp130 ΔAF/16683 Fab complex (EMD-73023 and 9YJG). Source data are provided with this paper. Regeneron materials described in this manuscript may be made available to qualified, academic, noncommercial researchers through a materials transfer agreement upon request at https://regeneron.envisionpharma.com/vt_regeneron/. For questions about how Regeneron shares materials, use the email address.

## Acknowledgments

We thank Bian Li, Kei Saotome, Johnathon Walls, Jennifer Cheng, and Harish Kumar Sura for discussion of the project, and Regeneron TP-Protein Development team for purification of the H4xH16683P2 antibody.

## Author contributions

Y.Z., M.W.S., W.C.O., P.E.S., and M.C.F. conceptualized the studies. Y.Z. prepared complexes, acquired cryo-EM data, processed data, and built and refined the atomic models. P.E.S., J.C., and J.J. purified the soluble protein reagents and performed the activity assays. T.R., L.L.M., and R. L. prepared full-length gp130 WT and mutants in detergent. A.S.I.C. interpreted the data and created the signaling models. M.W.S., W.C.O., and M.C.F. supervised the project. Y.Z. drafted the manuscript, which was edited and finalized with input from all authors.

## References

1. Poussin, K. et al. Biochemical and functional analyses of gp130 mutants unveil JAK1 as a novel therapeutic target in human inflammatory hepatocellular adenoma. Oncoimmunology 2, e27090 (2013).

2. Nault, J.C., Bioulac-Sage, P. & Zucman-Rossi, J. Hepatocellular benign tumors-from molecular classification to personalized clinical care. Gastroenterology 144, 888–902 (2013).

3. Bioulac-Sage, P. et al. Hepatocellular adenoma subtype classification using molecular markers and immunohistochemistry. Hepatology 46, 740–8 (2007).

4. Rebouissou, S. et al. Frequent in-frame somatic deletions activate gp130 in inflammatory hepatocellular tumours. Nature 457, 200–4 (2009).

5. Pilati, C. et al. Somatic mutations activating STAT3 in human inflammatory hepatocellular adenomas. J Exp Med 208, 1359–66 (2011).

6. Guichard, C. et al. Integrated analysis of somatic mutations and focal copy-number changes identifies key genes and pathways in hepatocellular carcinoma. Nat Genet 44, 694–8 (2012).

7. Amaddeo, G., Guichard, C., Imbeaud, S. & Zucman-Rossi, J. Next-generation sequencing identified new oncogenes and tumor suppressor genes in human hepatic tumors. Oncoimmunology 1, 1612–1613 (2012).

8. Schmidt-Arras, D. et al. Oncogenic deletion mutants of gp130 signal from intracellular compartments. J Cell Sci 127, 341–53 (2014).

9. Zhou, Y. et al. Structural insights into the assembly of gp130 family cytokine signaling complexes. Sci Adv 9, eade4395 (2023).

10. Zhou, Y. et al. Structures of complete extracellular assemblies of type I and type II Oncostatin M receptor complexes. Nat Commun 15, 9776 (2024).

11. Boulanger, M.J., Chow, D.C., Brevnova, E.E. & Garcia, K.C. Hexameric structure and assembly of the interleukin-6/IL-6 alpha-receptor/gp130 complex. Science 300, 2101–4 (2003).

12. Bravo, J., Staunton, D., Heath, J.K. & Jones, E.Y. Crystal structure of a cytokine-binding region of gp130. EMBO J 17, 1665–74 (1998).

13. Schutt, A. et al. gp130 activation is regulated by D2-D3 interdomain connectivity. Biochem J 450, 487–96 (2013).

14. Boulanger, M.J., Bankovich, A.J., Kortemme, T., Baker, D. & Garcia, K.C. Convergent mechanisms for recognition of divergent cytokines by the shared signaling receptor gp130. Mol Cell 12, 577–89 (2003).

15. Grebenciucova, E. & VanHaerents, S. Interleukin 6: at the interface of human health and disease. Front Immunol 14, 1255533 (2023).

16. Sommer, J. et al. Constitutively active mutant gp130 receptor protein from inflammatory hepatocellular adenoma is inhibited by an anti-gp130 antibody that specifically neutralizes interleukin 11 signaling. J Biol Chem 287, 13743–51 (2012).

17. Nandwani, N., Surana, P., Udgaonkar, J.B., Das, R. & Gosavi, S. Amino-acid composition after loop deletion drives domain swapping. Protein Sci 26, 1994–2002 (2017).

18. Nandwani, N. et al. A five-residue motif for the design of domain swapping in proteins. Nat Commun 10, 452 (2019).

19. Glassman, C.R. et al. Structure of a Janus kinase cytokine receptor complex reveals the basis for dimeric activation. Science 376, 163–169 (2022).

20. Caveney, N.A. et al. Structural basis of Janus kinase trans-activation. Cell Rep 42, 112201 (2023).

21. Lv, Y. et al. The JAK-STAT pathway: from structural biology to cytokine engineering. Signal Transduct Target Ther 9, 221 (2024).

22. Mohan, K. et al. Topological control of cytokine receptor signaling induces differential effects in hematopoiesis. Science 364(2019).

23. Vanamee, E.S., Lippner, G. & Faustman, D.L. Signal Amplification in Highly Ordered Networks Is Driven by Geometry. Cells 11(2022).

24. Kurth, I. et al. Importance of the membrane-proximal extracellular domains for activation of the signal transducer glycoprotein 130. J Immunol 164, 273–82 (2000).

25. Timmermann, A., Kuster, A., Kurth, I., Heinrich, P.C. & Muller-Newen, G. A functional role of the membrane-proximal extracellular domains of the signal transducer gp130 in heterodimerization with the leukemia inhibitory factor receptor. Eur J Biochem 269, 2716–26 (2002).

26. Wilson, S.C. et al. Organizing structural principles of the IL-17 ligand-receptor axis. Nature 609, 622–629 (2022).

27. Xu, Y. et al. Crystal structure of the entire ectodomain of gp130: insights into the molecular assembly of the tall cytokine receptor complexes. J Biol Chem 285, 21214–8 (2010).

28. Jumper, J. et al. Highly accurate protein structure prediction with AlphaFold. Nature 596, 583–589 (2021).

29. Usacheva, A. et al. Contribution of the Box 1 and Box 2 motifs of cytokine receptors to Jak1 association and activation. J Biol Chem 277, 48220–6 (2002).

30. Cairo, C.W. Signaling by committee: receptor clusters determine pathways of cellular activation. ACS Chem Biol 2, 652–5 (2007).

31. Westerfield, J.M. & Barrera, F.N. Membrane receptor activation mechanisms and transmembrane peptide tools to elucidate them. J Biol Chem 295, 1792–1814 (2020).

32. Chen, Z., Oh, D., Biswas, K.H., Zaidel-Bar, R. & Groves, J.T. Probing the effect of clustering on EphA2 receptor signaling efficiency by subcellular control of ligand-receptor mobility. Elife 10(2021).

33. Vanamee, E.S. & Faustman, D.L. The benefits of clustering in TNF receptor superfamily signaling. Front Immunol 14, 1225704 (2023).

34. Giese, B. et al. Dimerization of the cytokine receptors gp130 and LIFR analysed in single cells. J Cell Sci 118, 5129–40 (2005).

35. Tenhumberg, S. et al. gp130 dimerization in the absence of ligand: preformed cytokine receptor complexes. Biochem Biophys Res Commun 346, 649–57 (2006).

36. Rinis, N., Kuster, A., Schmitz-Van de Leur, H., Mohr, A. & Muller-Newen, G. Intracellular signaling prevents effective blockade of oncogenic gp130 mutants by neutralizing antibodies. Cell Commun Signal 12, 14 (2014).

37. Punjani, A., Rubinstein, J.L., Fleet, D.J. & Brubaker, M.A. cryoSPARC: algorithms for rapid unsupervised cryo-EM structure determination. Nat Methods 14, 290–296 (2017).

38. Pettersen, E.F. et al. UCSF Chimera--a visualization system for exploratory research and analysis. J Comput Chem 25, 1605–12 (2004).

39. Emsley, P., Lohkamp, B., Scott, W.G. & Cowtan, K. Features and development of Coot. Acta Crystallogr D Biol Crystallogr 66, 486–501 (2010).

40. Liebschner, D. et al. Macromolecular structure determination using X-rays, neutrons and electrons: recent developments in Phenix. Acta Crystallogr D Struct Biol 75, 861–877 (2019).

41. Williams, C.J. et al. MolProbity: More and better reference data for improved all-atom structure validation. Protein Sci 27, 293–315 (2018).

