## Supplemental_figures_tables for "Structural basis for constitutive activation of oncogenic gp130 mutants in human inflammatory hepatocellular adenomas"

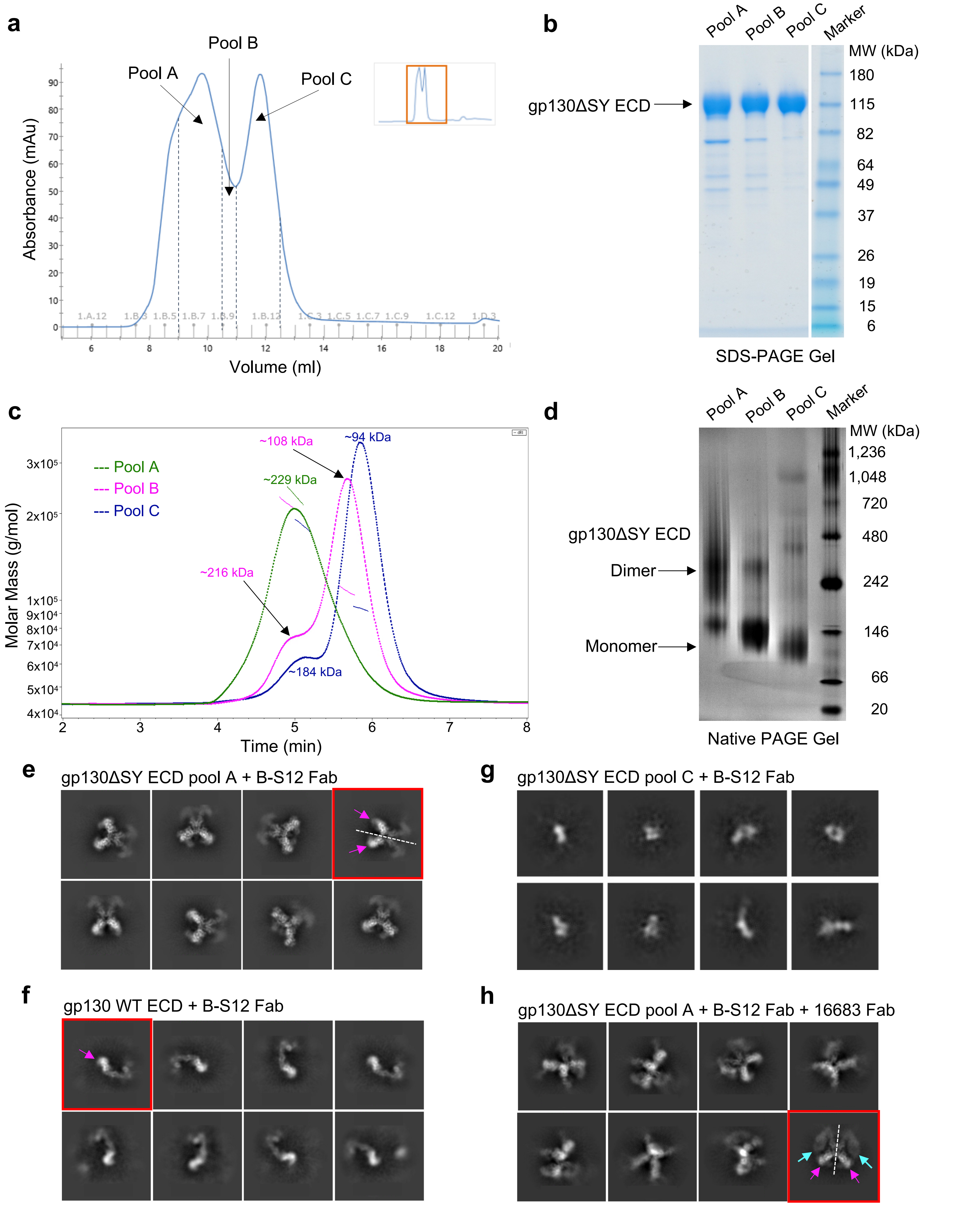


**Supplementary Figure 1. Purification and initial cryo-EM analysis of gp130 S187-Y190 deletion mutant extracellular domain (gp130 ΔSY ECD). a,** SEC profile of gp130 ΔSY ECD purification using a Superdex 200 Increase 10/300 GL column. Fractions under the two distinct peaks were collected as pool A and pool C, respectively. The fraction at the boundary was collected as pool B. **b,** SDS-PAGE gel of the three pooled samples in **a**. **c,** SEC-MALS analysis of the three pooled samples in **a**. **d,** Native PAGE gel of the three pooled samples in **a**. **e,** Representative 2D class averages of gp130 ΔSY ECD pool A sample in complex with B-S12 Fab. In the class highlighted by red square, the magenta arrow indicates B-S12 Fab while the dash line indicates symmetry axis of the complex. **f,** Representative 2D class averages of gp130 WT ECD protein in complex with B-S12 Fab. In the class highlighted by red square, the magenta arrow indicates B-S12 Fab. **g,** Representative 2D class averages of gp130 ΔSY ECD pool C sample in complex with B-S12 Fab. **h,** Representative 2D class averages of gp130 ΔSY ECD pool A sample in complex with B-S12 Fab and 16683 Fab. In the class highlighted in red square, the magenta arrow and cyan arrow indicate B-S12 Fab and 16683 Fab, respectively, while the dash line indicates symmetry axis of the complex.

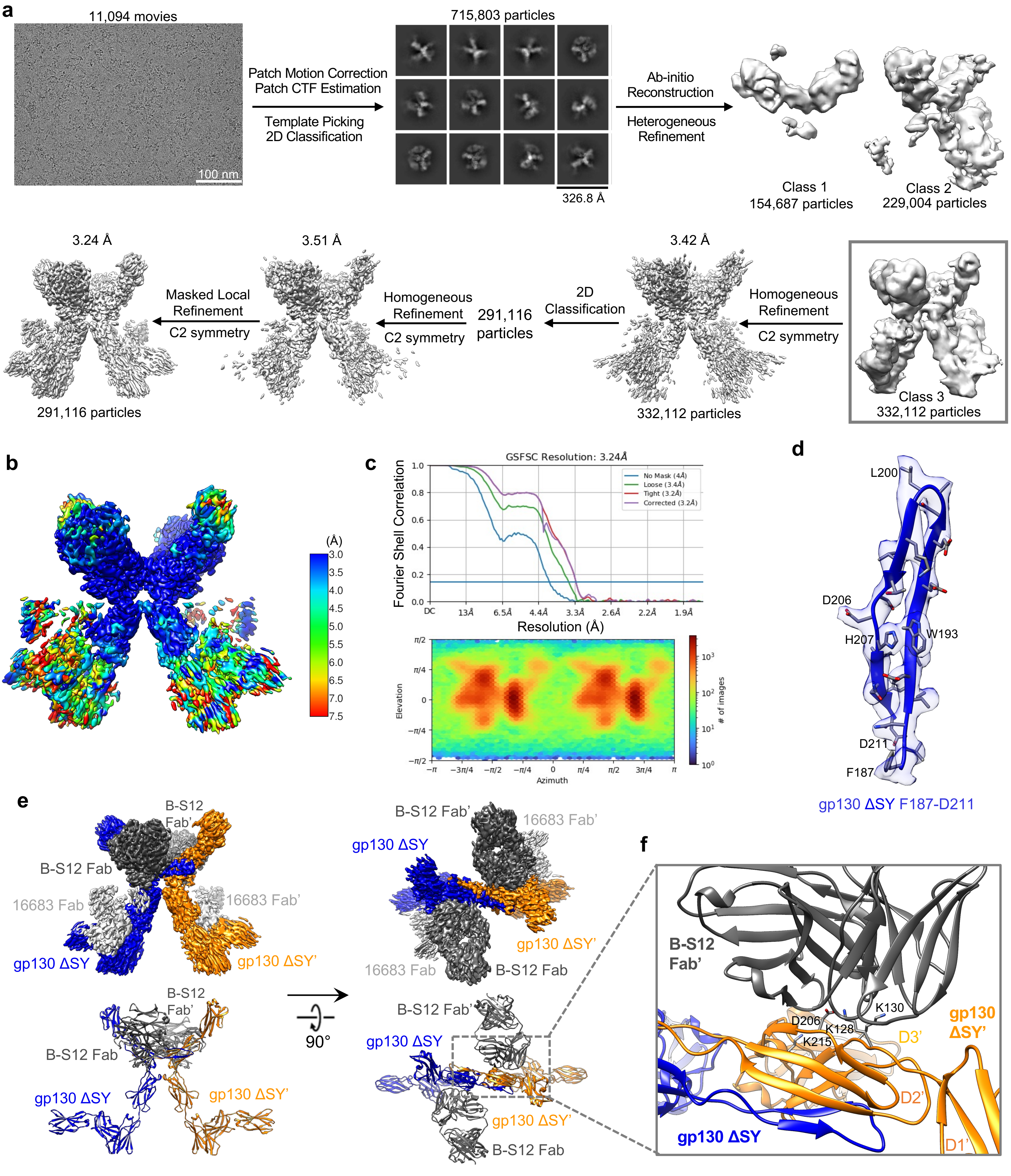


**Supplementary Figure 2. Cryo-EM structure of gp130 S187-Y190 deletion mutant extracellular domain (gp130 ΔSY ECD) in complex with B-S12 Fab and 16683 Fab. a,** Workflow for cryo-EM data processing. **b,** Local resolution estimation of the final cryo-EM map. **c,** FSC curve showing a global resolution of 3.24 Å based on the 0.143 gold standard threshold (top panel), and angular distribution plot (bottom panel). **d,** Representative cryo-EM density of the map. **e,** Side view and top-down view of cryo-EM density map (top) and atomic model (bottom) of the gp130 ΔSY ECD dimer in complex with B-S12 Fab and 16683 Fab. 16683 Fab was not modeled due to poor density. **f,** Zoomed in view of B-S12 binding interface showing that B-S12 Fab does not contribute to gp130 ΔSY ECD dimerization. Four charged residues in D2 of gp130 ΔSY that play critical roles in B-S12 binding, K128, K130, D206, and K215, are shown in stick representation.

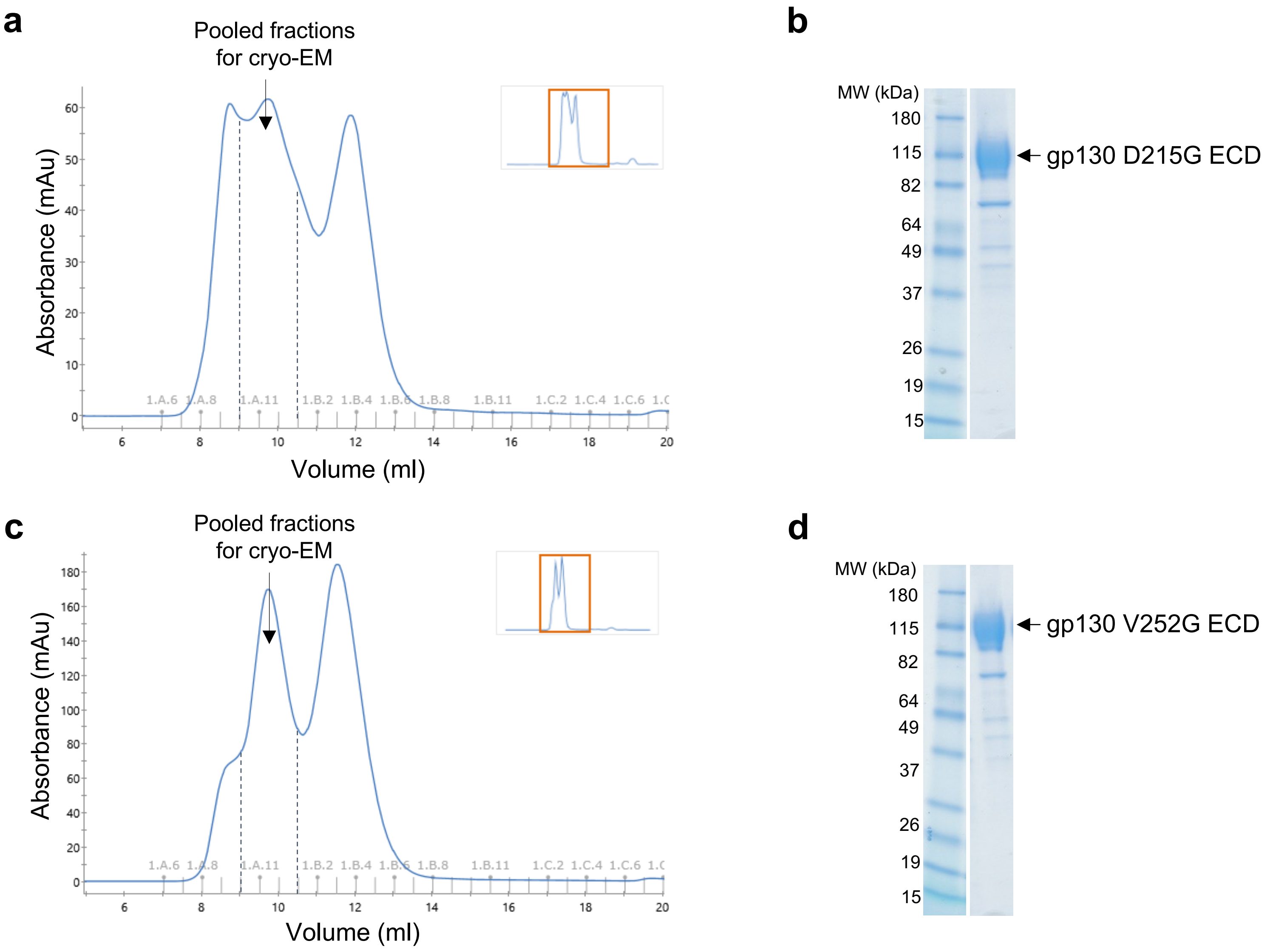


**Supplementary Figure 3. Purification of gp130 D215G mutant extracellular domain (gp130 D215G ECD) and gp130 V252G mutant extracellular domain (gp130 V252G ECD). a,** SEC profile of gp130 D215G ECD purification using a Superdex 200 Increase 10/300 GL column. The indicated fractions were pooled for cryo-EM analysis. **b,** SDS-PAGE gel of the pooled sample in **a**. **c,** SEC profile of gp130 V252G ECD purification using a Superdex 200 Increase 10/300 GL column. The indicated fractions were pooled for cryo-EM analysis. **b,** SDS-PAGE gel of the pooled sample in **c**.

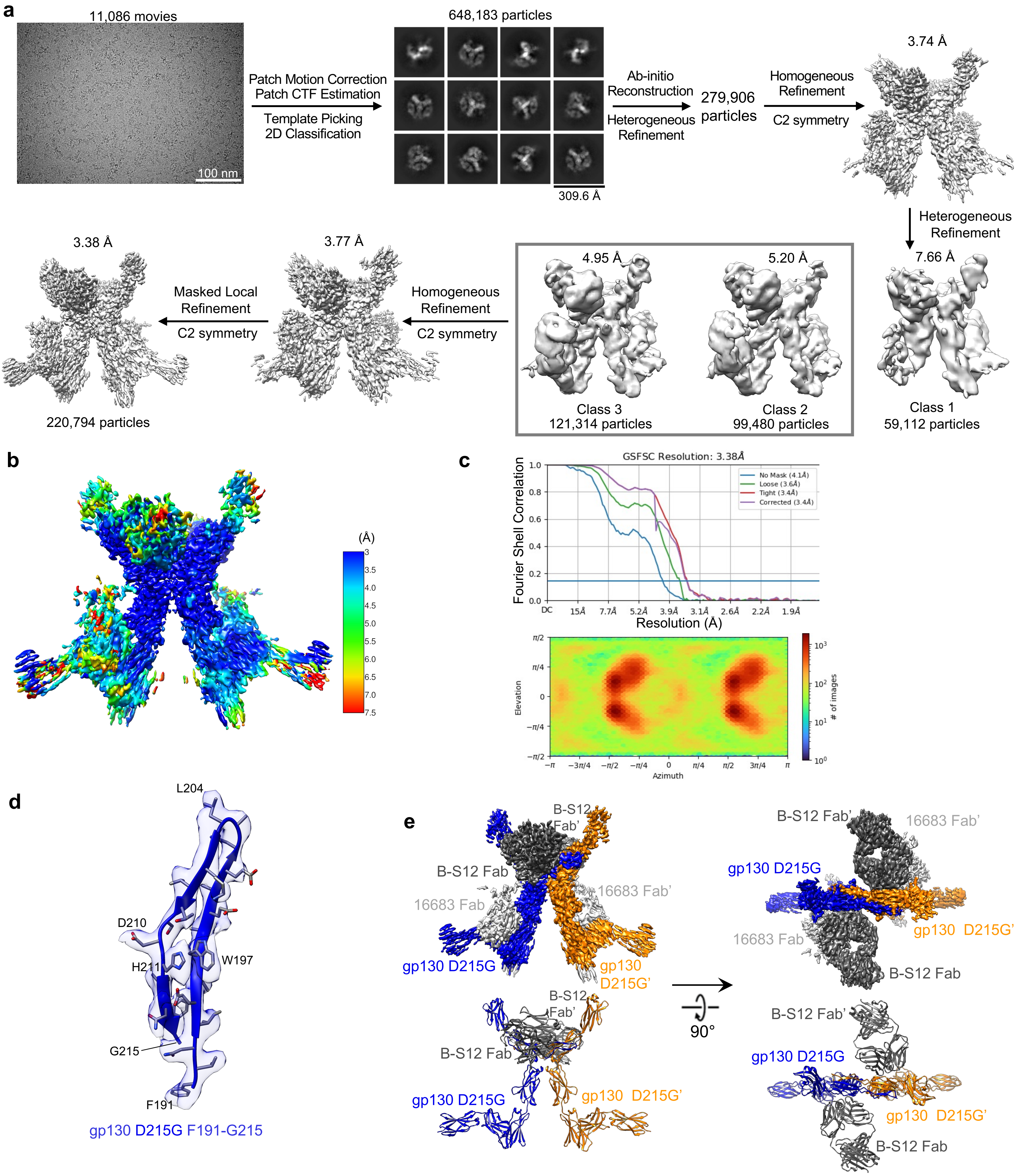


**Supplementary Figure 4. Cryo-EM structure of gp130 D215G mutant extracellular domain (gp130 D215G ECD) in complex with B-S12 Fab and 16683 Fab. a,** Workflow for cryo-EM data processing. **b,** Local resolution estimation of the final cryo-EM map. **c,** FSC curve showing a global resolution of 3.38 Å based on the 0.143 gold standard threshold (top panel), and angular distribution plot (bottom panel). **d,** Representative cryo-EM density of the map. **e,** Side view and top-down view of cryo-EM density map (top) and atomic model (bottom) of the gp130 D215G ECD dimer in complex with B-S12 Fab and 16683 Fab. 16683 Fab was not modeled due to poor density.

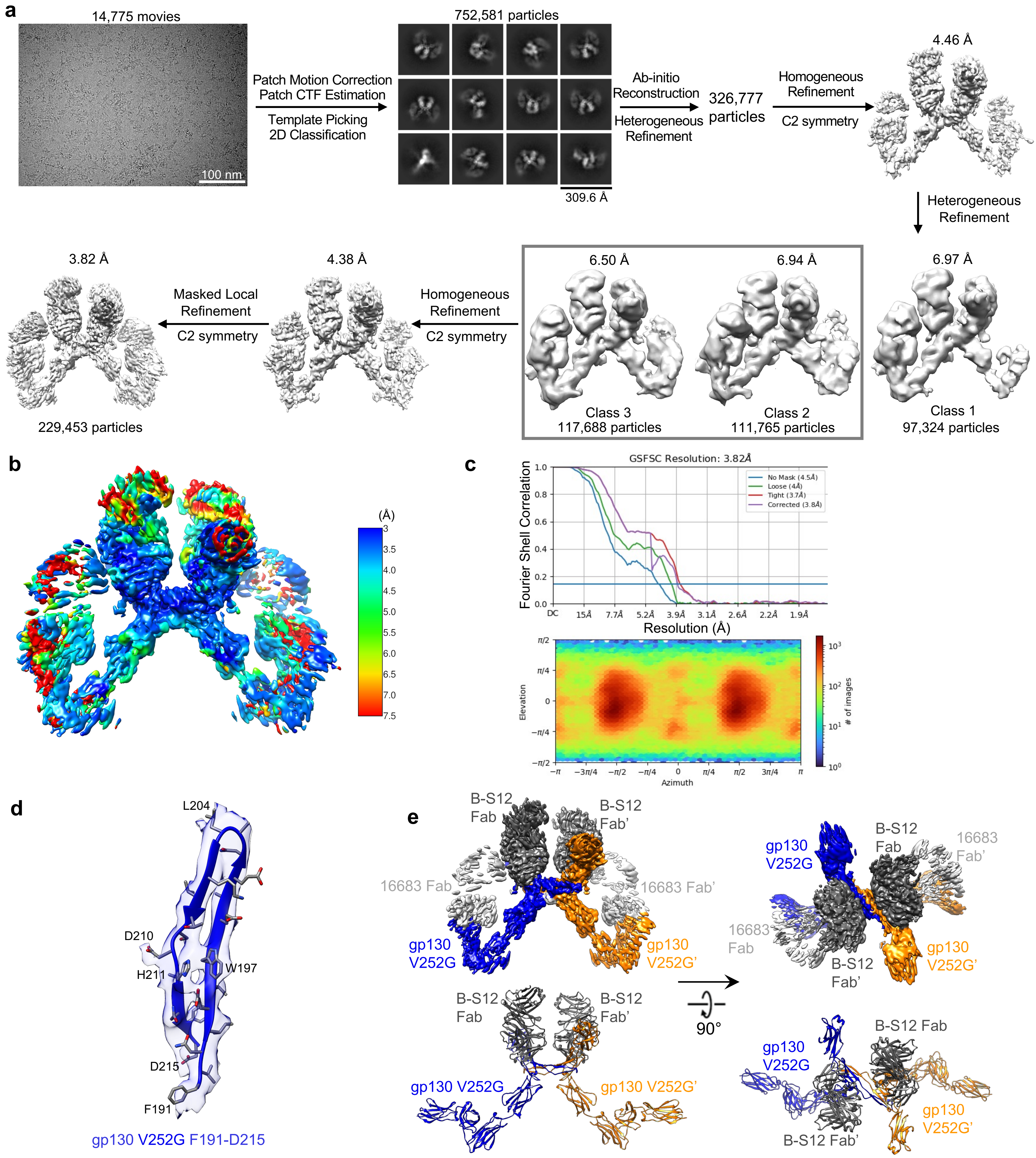


**Supplementary Figure 5. Cryo-EM structure of gp130 V252G mutant extracellular domain (gp130 V252G ECD) in complex with B-S12 Fab and 16683 Fab. a,** Workflow for cryo-EM data processing. **b,** Local resolution estimation of the final cryo-EM map. **c,** FSC curve showing a global resolution of 3.82 Å based on the 0.143 gold standard threshold (top panel), and angular distribution plot (bottom panel). **d,** Representative cryo-EM density of the map. **e,** Side view and top-down view of cryo-EM density map (top) and atomic model (bottom) of the gp130 V252G ECD dimer in complex with B-S12 Fab and 16683 Fab. 16683 Fab was not modeled due to poor density.

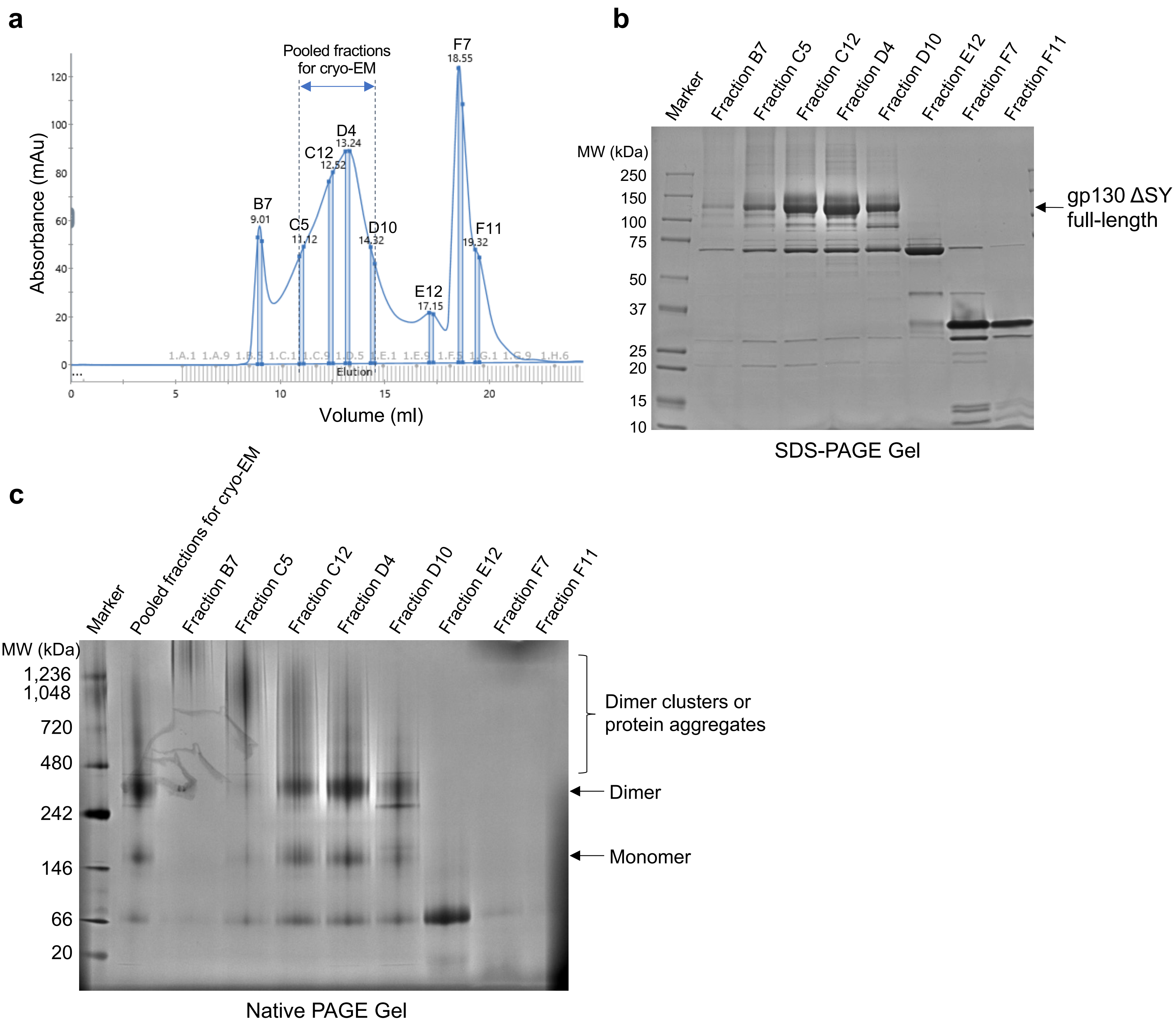


**Supplementary Figure 6. Purification of full-length gp130 S187-Y190 deletion mutant (gp130 ΔSY full-length) in detergent. a,** SEC profile of gp130 ΔSY full-length purification using a Superose 6 Increase 10/300 GL column. Fractions C5 to D10 were pooled for cryo-EM analysis. **b,** SDS-PAGE gel of different fraction samples in **a**. **c,** Native PAGE gel of the different fraction samples and pooled samples in **a**.

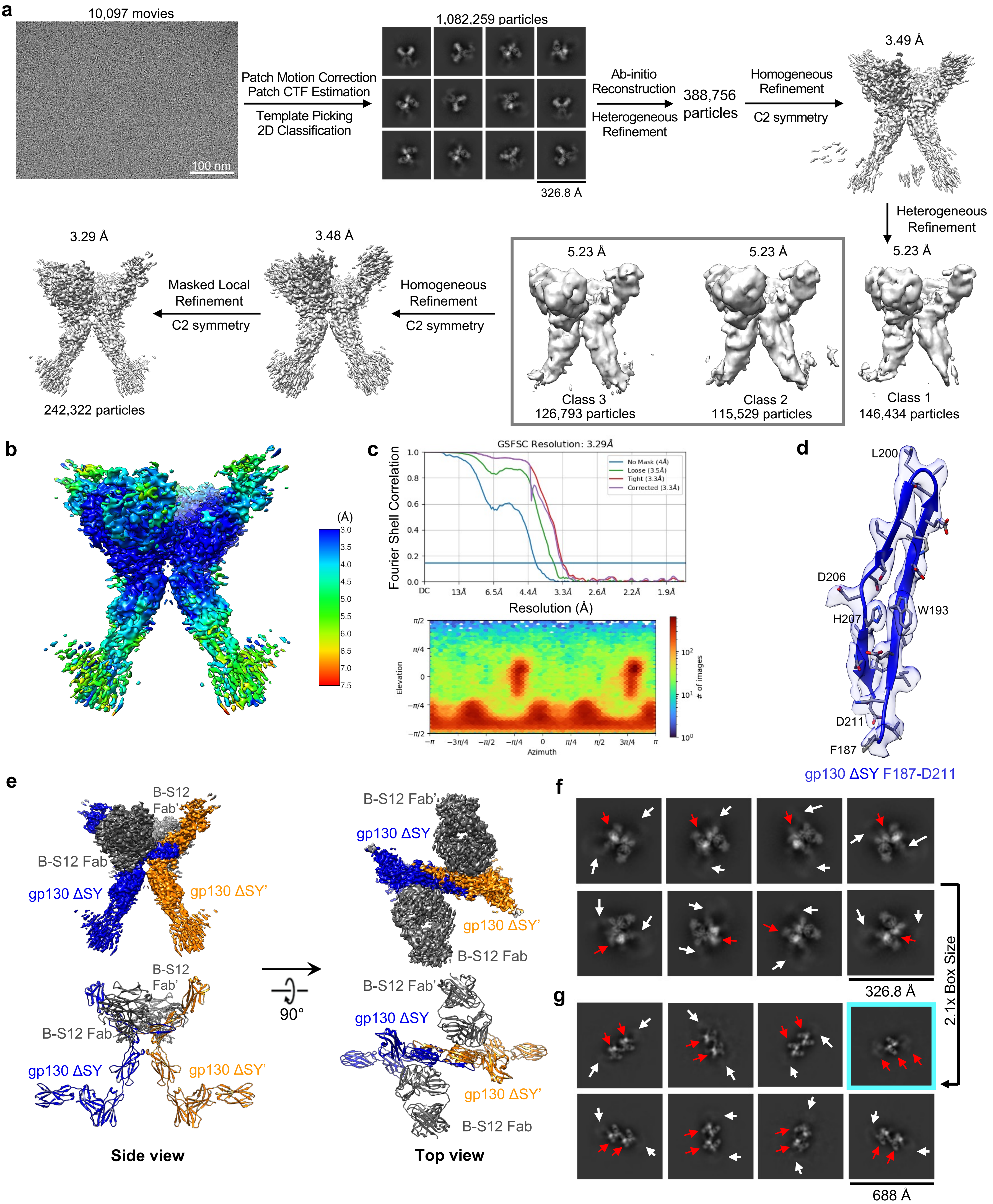


**Supplementary Figure 7. Cryo-EM structure of individual dimer of full-length gp130 S187-Y190 deletion mutant (gp130 ΔSY full-length) in complex with B-S12 Fab. a,** Workflow for cryo-EM data processing. **b,** Local resolution estimation of the final cryo-EM map. **c,** FSC curve showing a global resolution of 3.29 Å based on the 0.143 gold standard threshold (top panel), and angular distribution plot (bottom panel). **d,** Representative cryo-EM density of the map. **e,** Side view and top-down view of cryo-EM density map (top) and atomic model (bottom) of the gp130 ΔSY full-length dimer in complex with B-S12 Fab. **f,** Representative 2D class averages of gp130 ΔSY full-length/B-S12 Fab complex particles extracted with 326.8 Å box size at top-down/bottom-up view. The red arrow indicates gp130 ΔSY full-length dimer, and the white arrows indicate fuzzy density attached to both sides of the dimer. **g,** Representative 2D class averages of gp130 ΔSY full-length/B-S12 Fab complex particles extracted with 688 Å box size at top-down/bottom-up view. The red arrows indicate gp130 ΔSY full-length dimers in the dimer cluster, and the white arrows indicate fuzzy density attached to both sides of the dimer cluster. The class highlighted by cyan box shows a trimer of dimer.

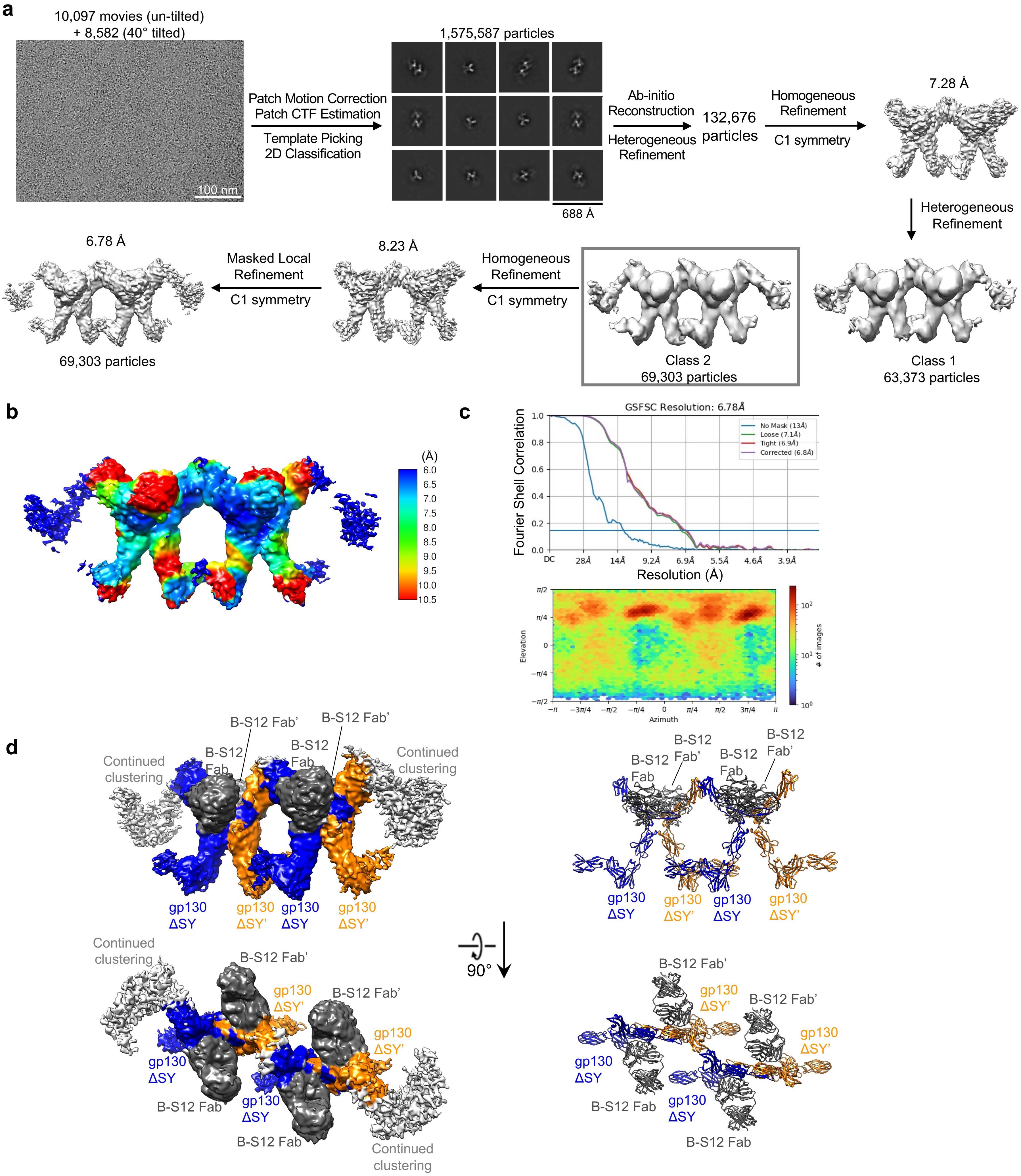


**Supplementary Figure 8. Cryo-EM structure of dimer cluster of full-length gp130 S187-Y190 deletion mutant (gp130 ΔSY full-length) in complex with B-S12 Fab. a,** Workflow for cryo-EM data processing. **b,** Local resolution estimation of the final cryo-EM map. **c,** FSC curve showing a global resolution of 6.78 Å based on the 0.143 gold standard threshold (top panel), and angular distribution plot (bottom panel). **d,** Side view and top-down view of cryo-EM density map (left) and atomic model (right) of the gp130 ΔSY full-length dimer cluster in complex with B-S12 Fab. The fragmented density attached to two sides of the dimer cluster likely indicates continued clustering of the dimer.

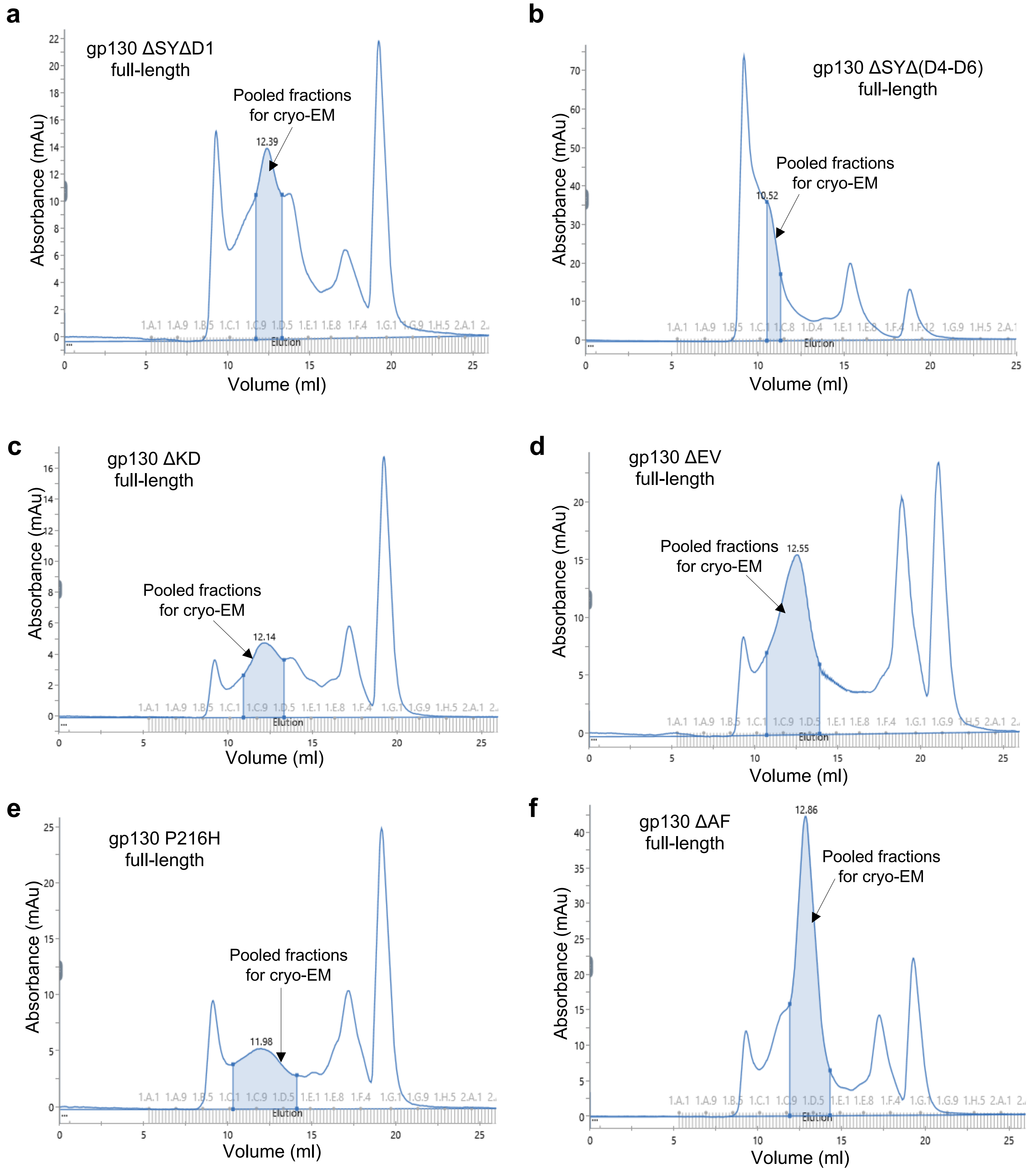


**Supplementary Figure 9. Purification of full-length gp130 mutants in detergent.** SEC profiles for purification of full-length gp130 S187-Y190 deletion mutant with additional D1 deletion (gp130 ΔSYΔD1) (**a**), gp130 S187-Y190 deletion mutant with additional D4-D6 deletion (gp130 ΔSYΔ(D4-D6)) (**b**), gp130 K173-D177 deletion mutant (gp130 ΔKD) (**c**), gp130 E195-V196 deletion mutant (gp130 ΔEV) (**d**), gp130 P216H mutant (**e**), and gp130 A418-F421 deletion mutant (gp130 ΔAF) (**f**) . Full-length gp130 ΔSYΔ(D4-D6) was purified using a Superdex 200 Increase 10/300 GL column while other proteins were purified using a Superose 6 Increase 10/300 GL column. Pooled fractions for cryo-EM studies are indicated.

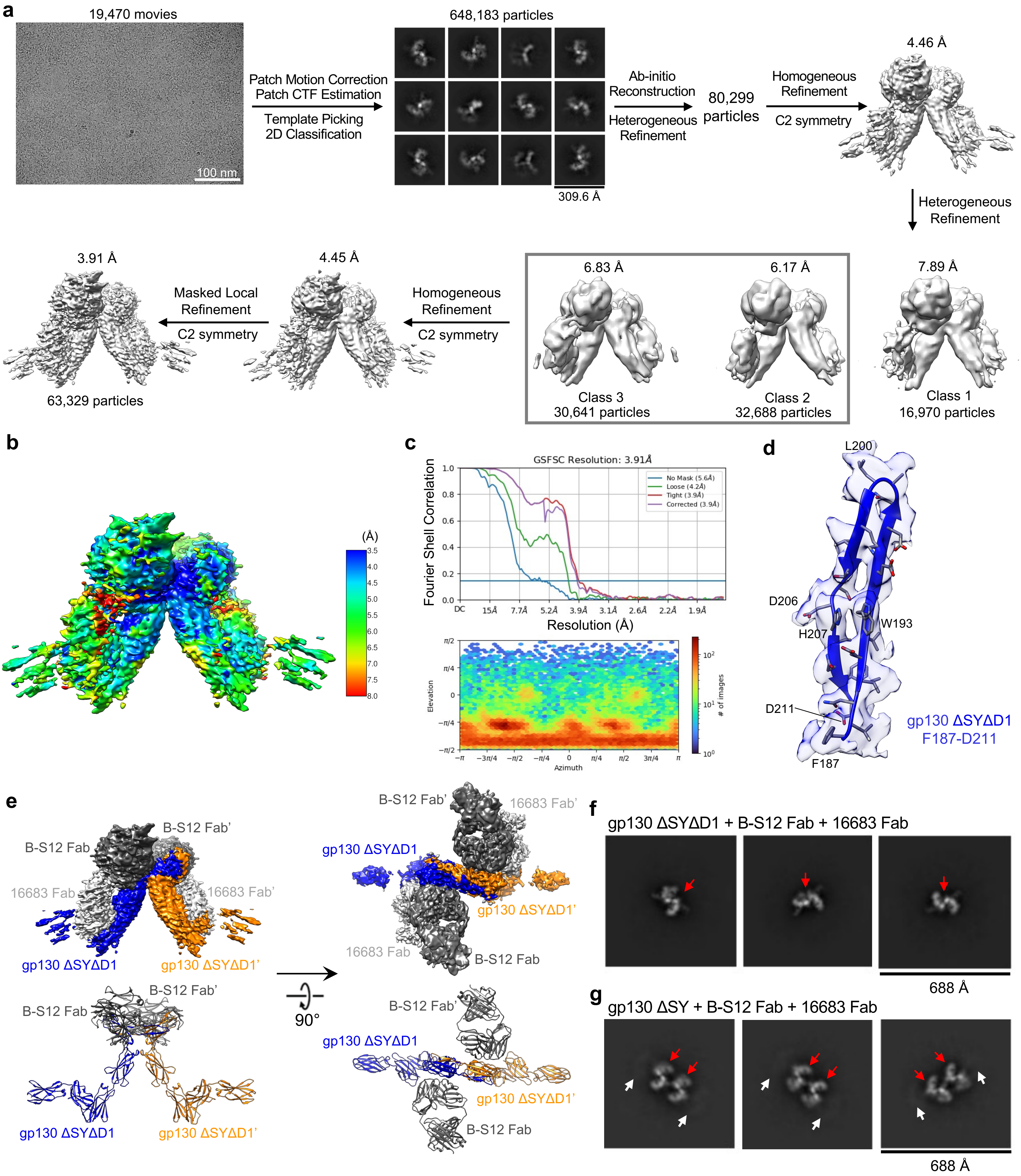


**Supplementary Figure 10. Cryo-EM structure of full-length gp130 S187-Y190 deletion mutant with additional D1 deletion (gp130 ΔSYΔD1 full-length) in complex with B-S12 Fab and 16683 Fab. a,** Workflow for cryo-EM data processing. **b,** Local resolution estimation of the final cryo-EM map. **c,** FSC curve showing a global resolution of 3.91 Å based on the 0.143 gold standard threshold (top panel), and angular distribution plot (bottom panel). **d,** Representative cryo-EM density of the map. **e,** Side view and top-down view of cryo-EM density map (top) and atomic model (bottom) of the gp130 ΔSYΔD1 full-length dimer in complex with B-S12 Fab and 16683 Fab. 16683 Fab was not modeled due to poor density. **f,** Representative 2D class averages of gp130 ΔSYΔD1 full-length/B-S12 Fab/16683 Fab complex particles extracted with 688 Å box size at top-down/bottom-up view. The red arrow indicates gp130 ΔSYΔD1 full-length dimer. **g,** Representative 2D class averages of gp130 ΔSY full-length/B-S12 Fab/16683 Fab complex particles extracted with 688 Å box size at top-down/bottom-up view. The red arrows indicate gp130 ΔSY full-length dimers in the dimer cluster, and the white arrows indicate fuzzy density attached to both sides of the dimer cluster.

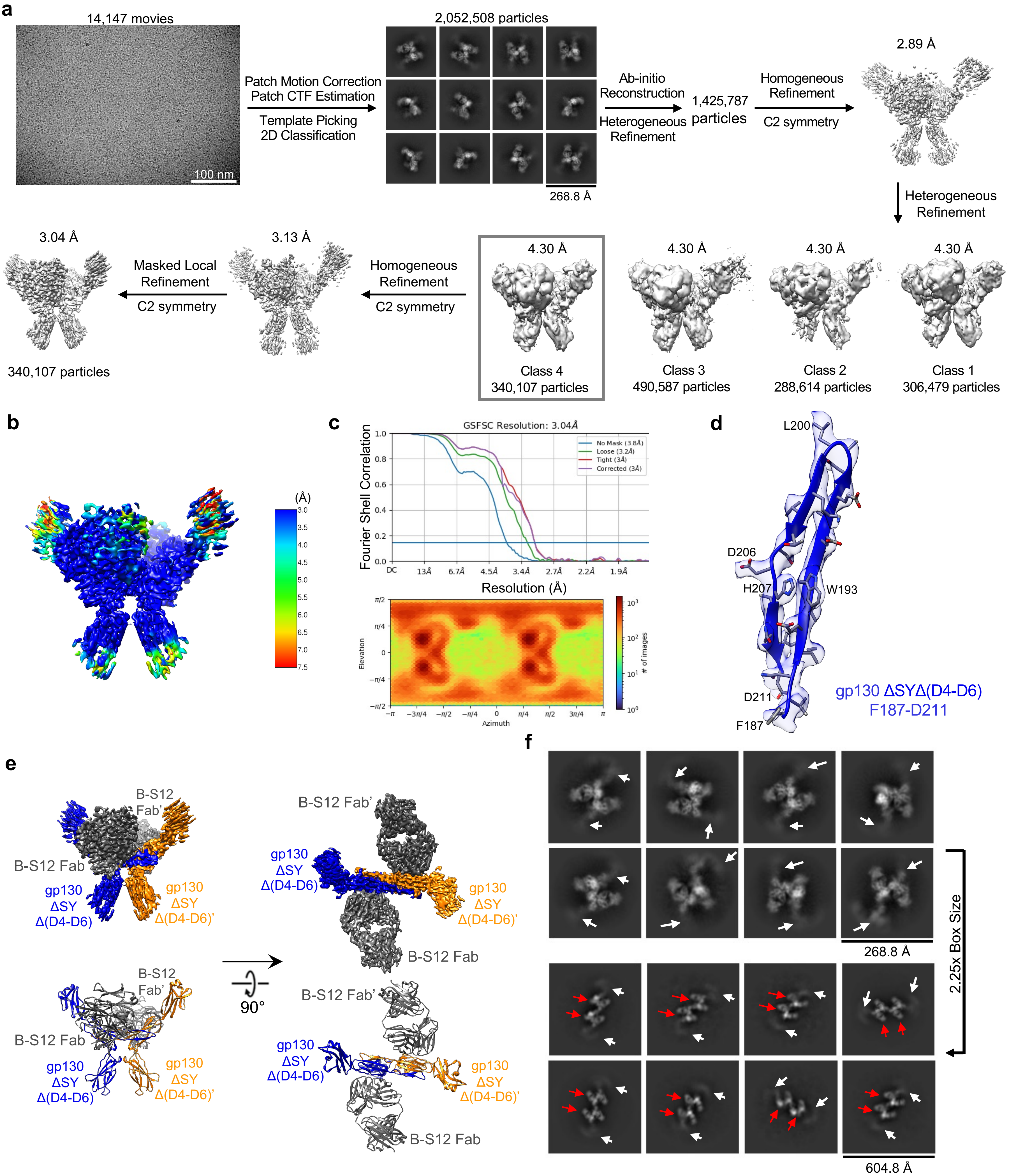


**Supplementary Figure 11. Cryo-EM structure of individual dimer of full-length gp130 S187-Y190 deletion mutant with additional D4-D6 deletion (gp130 ΔSYΔ(D4-D6) full-length) in complex with B-S12 Fab. a,** Workflow for cryo-EM data processing. **b,** Local resolution estimation of the final cryo-EM map. **c,** FSC curve showing a global resolution of 3.04 Å based on the 0.143 gold standard threshold (top panel), and angular distribution plot (bottom panel). **d,** Representative cryo-EM density of the map. **e,** Side view and top-down view of cryo-EM density map (top) and atomic model (bottom) of the gp130 ΔSYΔ(D4-D6) full-length dimer in complex with B-S12 Fab. **f,** Representative 2D class averages of gp130 ΔSYΔ(D4-D6) full-length/B-S12 Fab complex particles extracted with 268.8 Å box size at top-down/bottom-up view. The red arrow indicates gp130 ΔSY Δ(D4-D6) full-length dimer, and the white arrows indicate fuzzy density attached to both sides of the dimer. **g,** Representative 2D class averages of gp130 ΔSYΔ(D4-D6) full-length/B-S12 Fab complex particles extracted with 604.8 Å box size at top-down/bottom-up view. The red arrows indicate gp130 ΔSYΔ(D4-D6) full-length dimers in the dimer cluster, and the white arrows indicate fuzzy density attached to both sides of the dimer cluster.

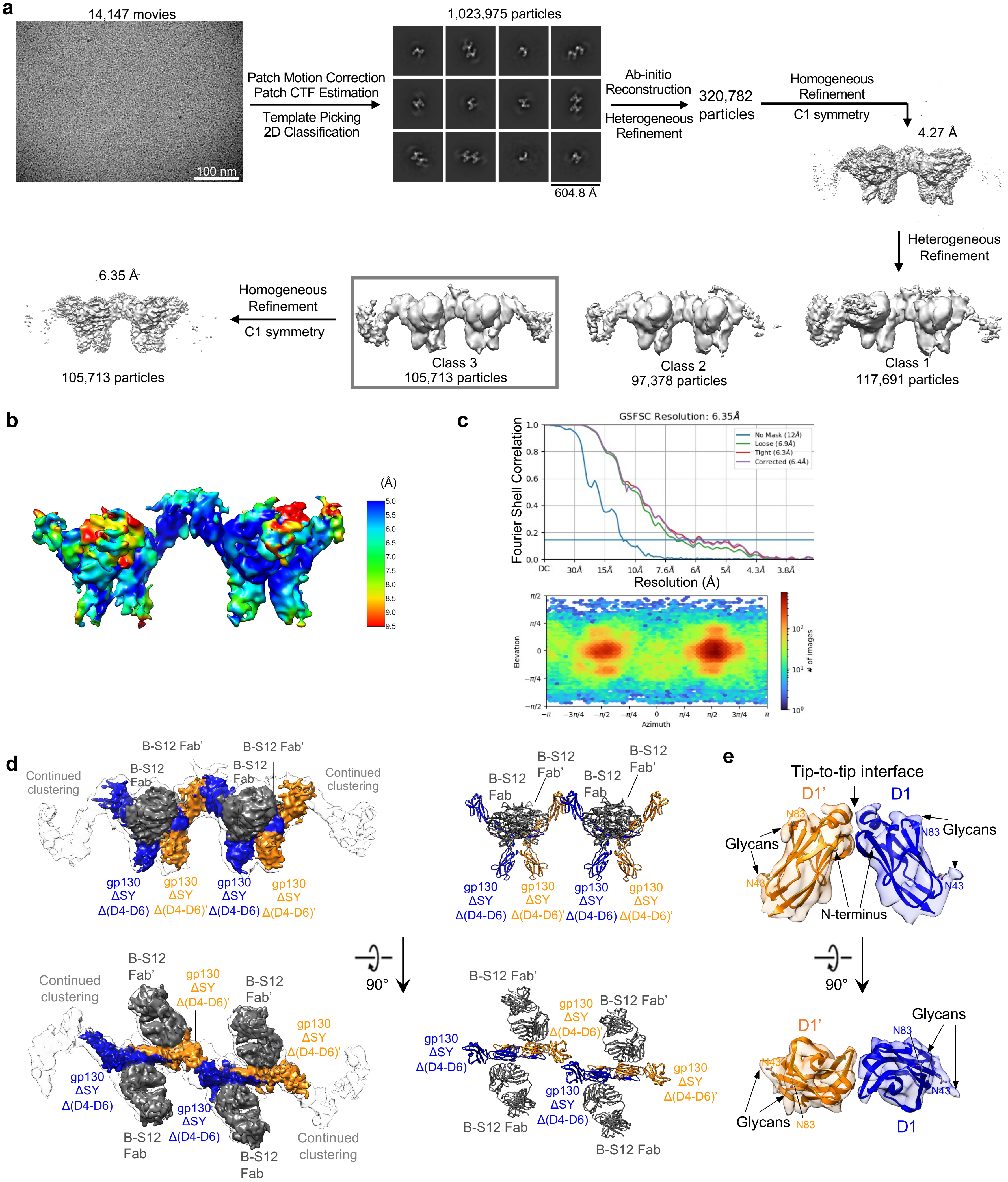


**Supplementary Figure 12. Cryo-EM structure of dimer cluster of full-length gp130 S187-Y190 deletion mutant with additional D4-D6 deletion (gp130 ΔSYΔ(D4-D6) full-length) in complex with B-S12 Fab. a,** Workflow for cryo-EM data processing. **b,** Local resolution estimation of the final cryo-EM map. **c,** FSC curve showing a global resolution of 6.35 Å based on the 0.143 gold standard threshold (top panel), and angular distribution plot (bottom panel). **d,** Side view and top-down view of cryo-EM density map (left) and atomic model (right) of the gp130 ΔSYΔ(D4-D6) full-length/B-S12 Fab complex. The colored final map is docked in the transparent intermediate map showing clear continued clustering at both sides. **e,** Zoomed-in views of the tip-to-tip interactions of D1 in the dimer cluster shown in **d**. The model docked in transparent density is shown at both side view and top-down view. The glycan density at N43 and N83 confirms the accuracy of the model docking.

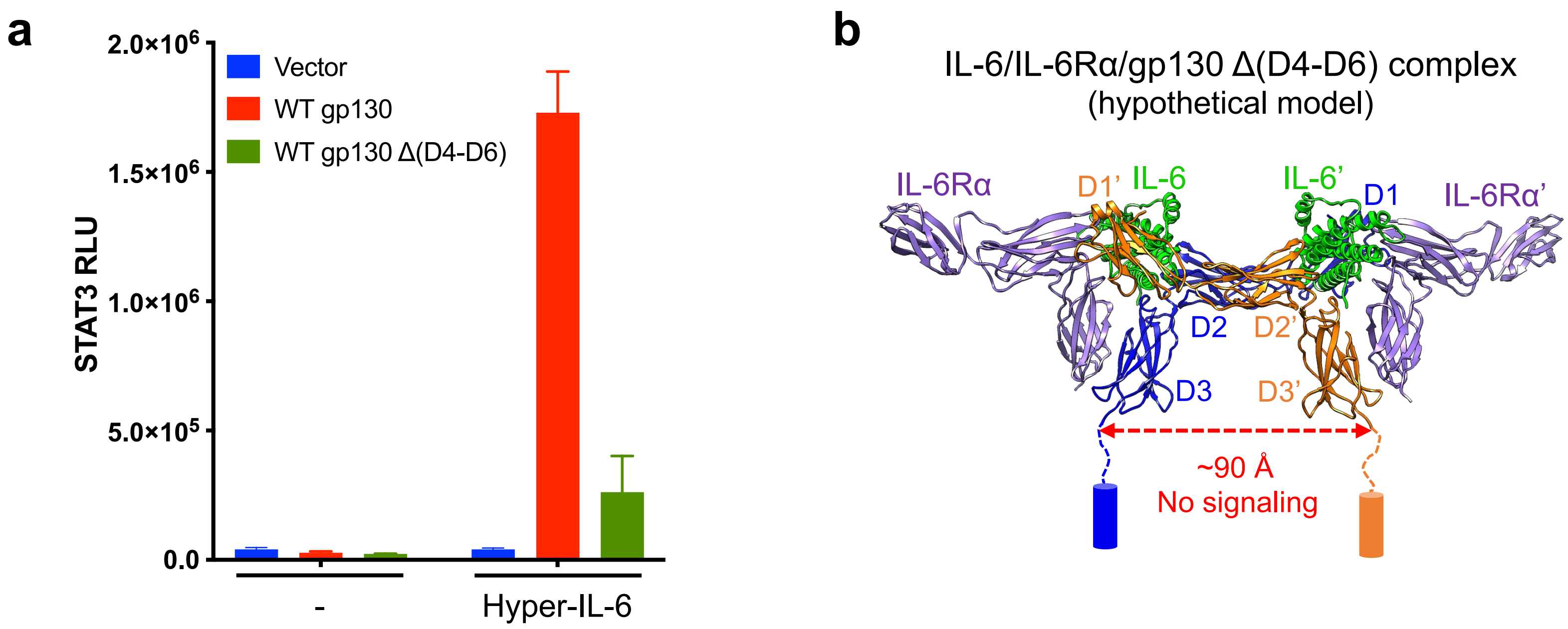


**Supplementary Figure 13. The FNIII domains (D4-D6) of WT gp130 are essential for IL-6-induced gp130 activation.**

**a** STAT3-luciferase reporter assay was performed in 293T cells after transiently overexpressing gp130 WT or gp130 Δ(D4-D6), with or without treatment by 10 nM Hyper-IL-6 (a fusion protein of IL-6 and its soluble receptor alpha subunit IL-6Rα). The data shown are representative of 2 independent experiments. RLU: relative luminescence units.

**b** Hypothetical model of IL-6/IL-6Rα/gp130 Δ(D4-D6) complex. The distance between the two juxtamembrane D3 domains is estimated. Cylinders represent TM domains.

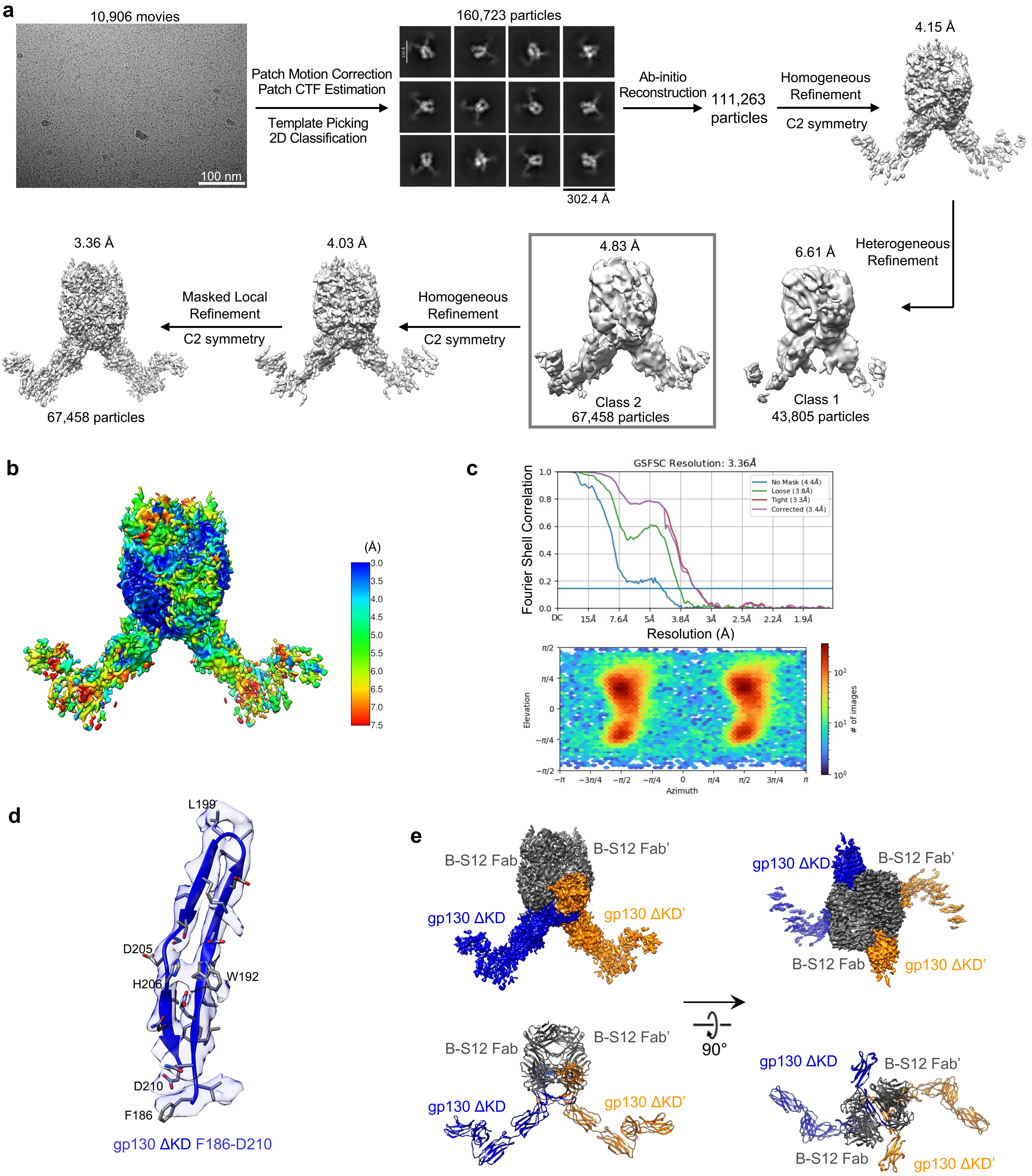


**Supplementary Figure 14. Cryo-EM structure of individual dimer of full-length gp130 K173-D177 deletion mutant (gp130 ΔKD full-length) in complex with B-S12 Fab. a,** Workflow for cryo-EM data processing. **b,** Local resolution estimation of the final cryo-EM map. **c,** FSC curve showing a global resolution of 3.36 Å based on the 0.143 gold standard threshold (top panel), and angular distribution plot (bottom panel). **d,** Representative cryo-EM density of the map. **e,** Side view and top-down view of cryo-EM density map (top) and atomic model (bottom) of the gp130 ΔKD full-length dimer in complex with B-S12 Fab.

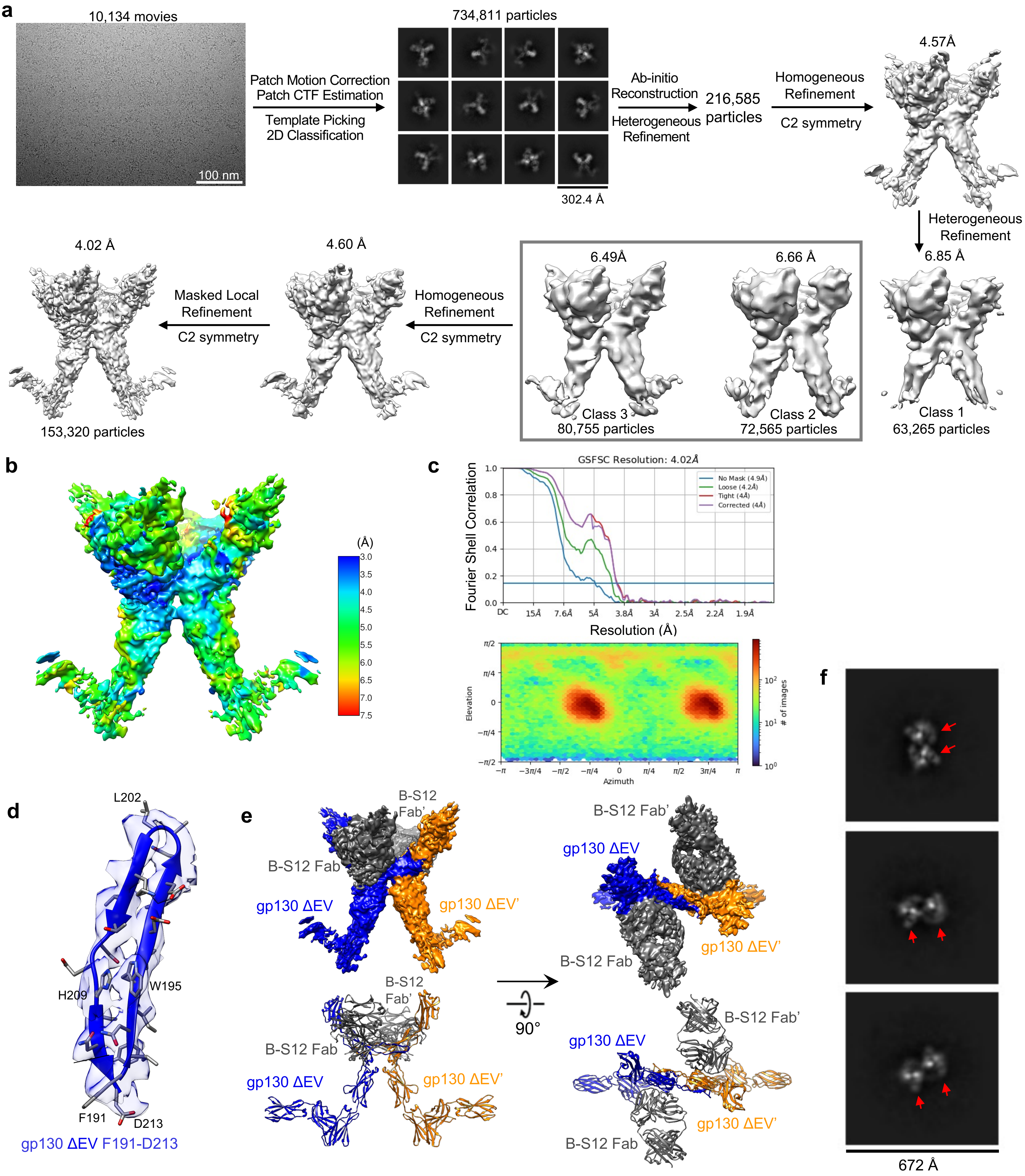


**Supplementary Figure 15. Cryo-EM structure of individual dimer of full-length gp130 E195-V196 deletion mutant (gp130 ΔEV full-length) in complex with B-S12 Fab. a,** Workflow for cryo-EM data processing. **b,** Local resolution estimation of the final cryo-EM map. **c,** FSC curve showing a global resolution of 4.02 Å based on the 0.143 gold standard threshold (top panel), and angular distribution plot (bottom panel). **d,** Representative cryo-EM density of the map. **e,** Side view and top-down view of cryo-EM density map (top) and atomic model (bottom) of the gp130 ΔEV full-length dimer in complex with B-S12 Fab. **f,** Representative 2D class averages of gp130 ΔEV full-length/B-S12 Fab complex particles extracted with 672 Å box size at top-down/bottom-up view. The red arrows indicate gp130 ΔEV full-length dimers in the dimer cluster.

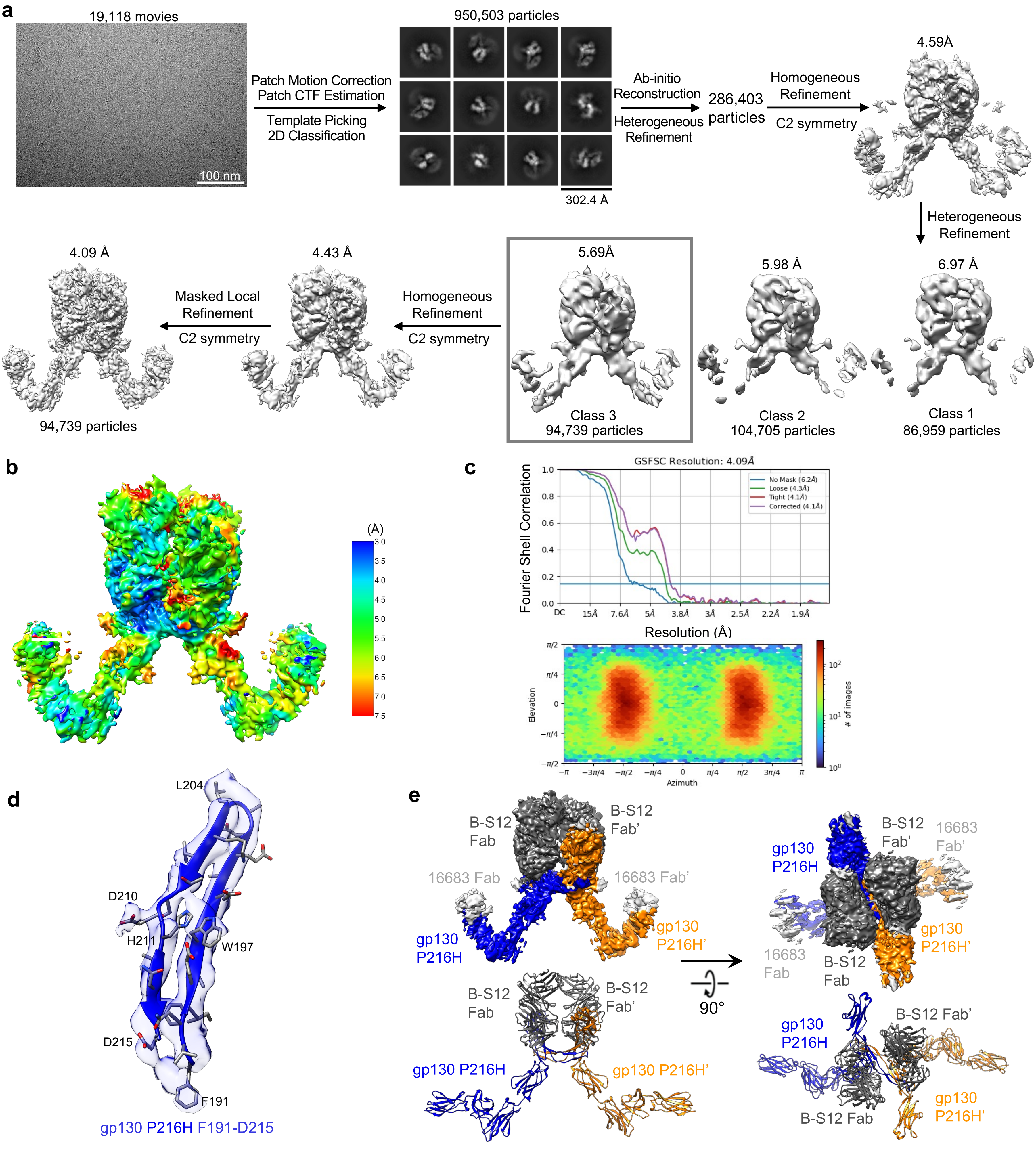


**Supplementary Figure 16. Cryo-EM structure of individual dimer of full-length gp130 P216H mutant (gp130 P216H full-length) in complex with B-S12 Fab and 16683 Fab. a,** Workflow for cryo-EM data processing. **b,** Local resolution estimation of the final cryo-EM map. **c,** FSC curve showing a global resolution of 4.09 Å based on the 0.143 gold standard threshold (top panel), and angular distribution plot (bottom panel). **d,** Representative cryo-EM density of the map. **e,** Side view and top-down view of cryo-EM density map (top) and atomic model (bottom) of the gp130 P216H full-length dimer in complex with B-S12 Fab and 16683 Fab. 16683 Fab was not modeled due to poor density.

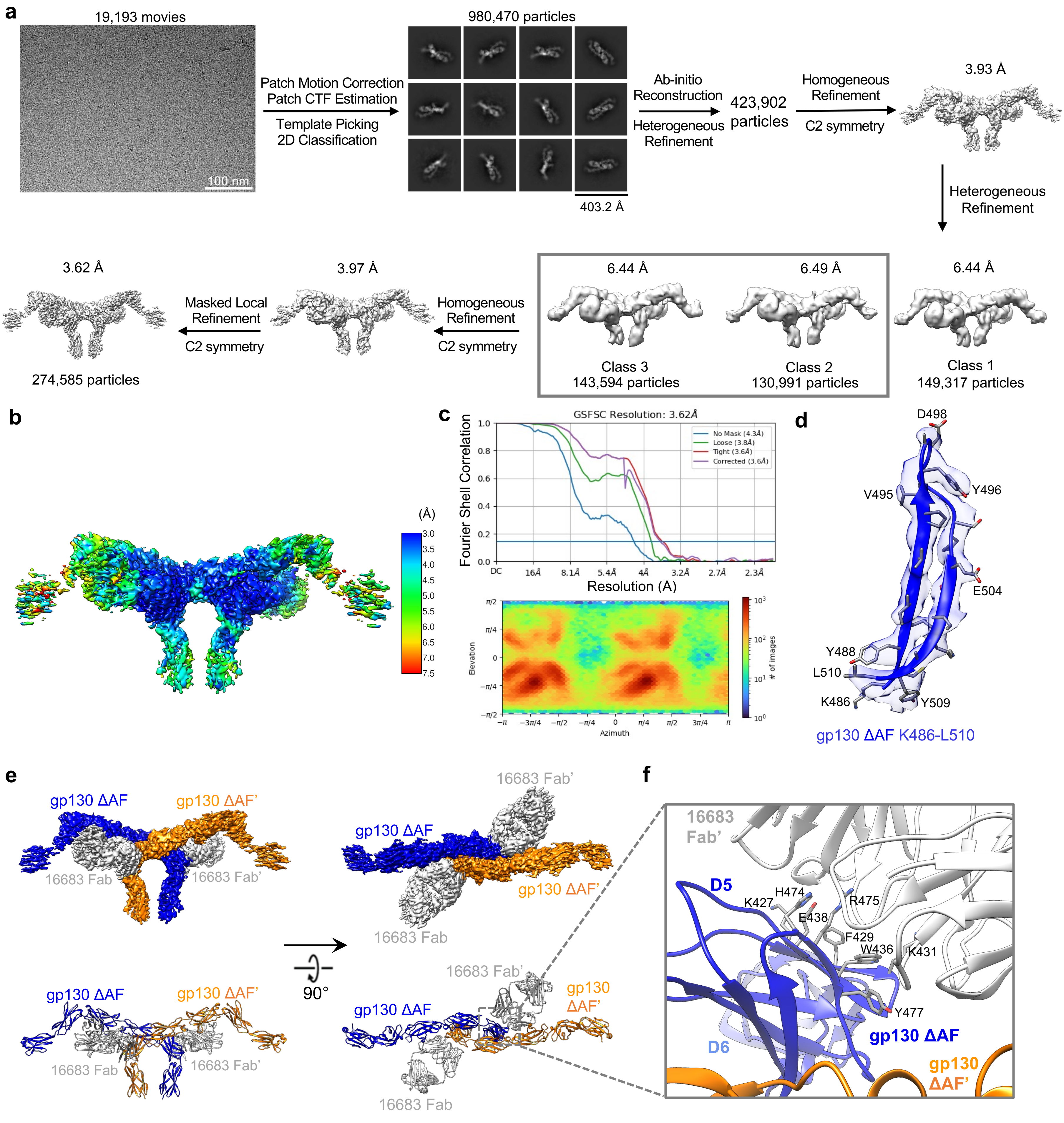


**Supplementary Figure 17. Cryo-EM structure of individual dimer of full-length gp130 A418-F421 deletion mutant (gp130 ΔAF full-length) in complex with 16683 Fab. a,** Workflow for cryo-EM data processing. **b,** Local resolution estimation of the final cryo-EM map. **c,** FSC curve showing a global resolution of 3.62 Å based on the 0.143 gold standard threshold (top panel), and angular distribution plot (bottom panel). **d,** Representative cryo-EM density of the map. **e,** Side view and top-down view of cryo-EM density map (top) and atomic model (bottom) of the gp130 ΔAF full-length dimer in complex with 16683 Fab. **f,** Zoomed in view of 16683 binding interface showing that 16683 Fab does not contribute to gp130 ΔAF dimerization. The residues in D5 of gp130 ΔAF that play critical roles in 16683 binding are shown in stick representation.

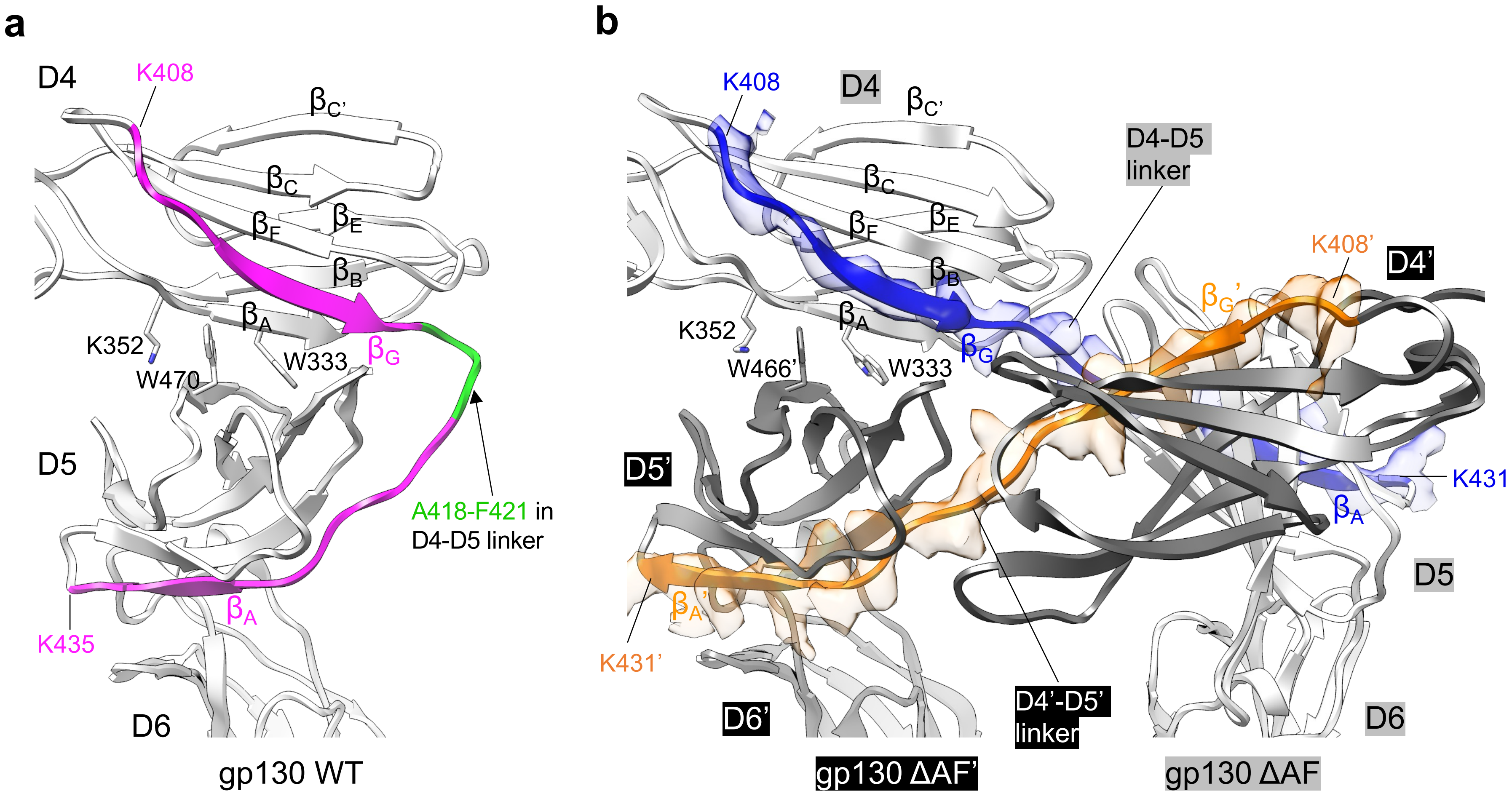


**Supplementary Figure 18. The acutely bent conformation of D4D5 observed in WT gp130 is retained in the gp130 ΔAF dimer through the interaction of D4 from one monomer (gp130 ΔAF) and D5' from the other monomer (gp130 ΔAF') following domain swapping between the two monomers.**

**a** WT gp130 has an acutely bent (~80°) conformation of D4D5. Three residues critical for maintaining this conformation, W333, K352, and W470, are shown in stick representation. D4 $\beta_G$, D5 $\beta_A$, and the D4D5 linker are colored in magenta. The deleted region (A418-F421) in gp130 ΔAF mutant is colored in green.

**b** gp130 ΔAF mutant dimer is shown in the same orientation as **a**. The two monomers, gp130 ΔAF and gp130 ΔAF', are colored in white and dark grey, respectively. D4 $\beta_G$, D5 $\beta_A$, and the D4D5 linker in transparent cryo-EM density of this region in the two monomers are colored in blue and orange, respectively. K352 and W333 from gp130 ΔAF D4, and W466' (W470 in WT gp130) from gp130 ΔAF' D5', that are important for maintaining the acutely bent conformation of D4D5', are highlighted in stick representation.

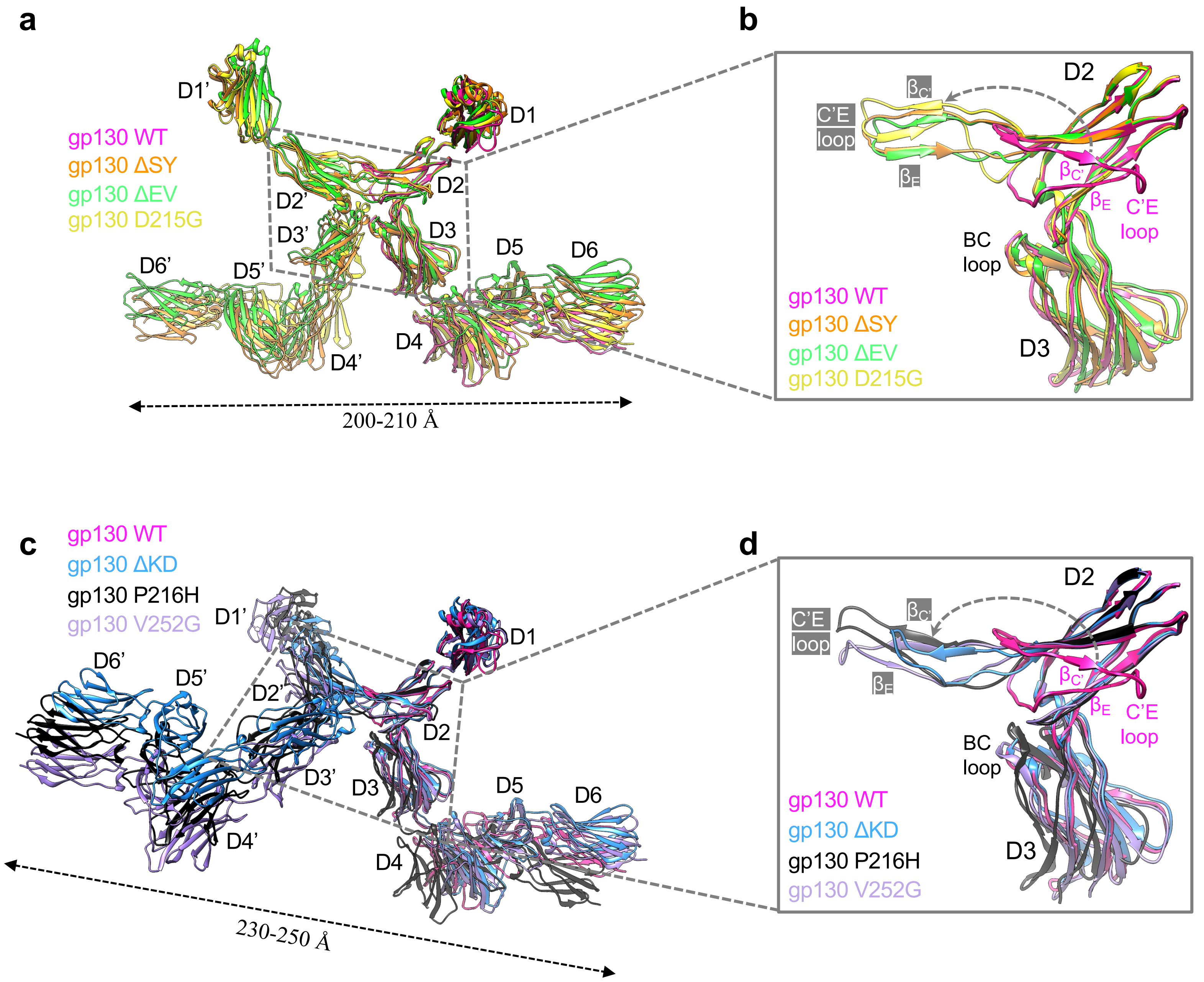


**Supplementary Figure 19. Structural comparison of WT gp130 and constitutively active gp130 mutants.**

**a** Overlay of WT gp130 (magenta), gp130 ΔSY (orange), gp130 ΔEV (green), and gp130 D215G (yellow).

**b** Zoomed-in view of D2D3 from structures in **a** showing distinct conformations of the βC'-C'E-βE modules in different mutants. Only one monomer in each mutant dimer is shown for better visibility.

**c** Overlay of WT gp130 (magenta), gp130 ΔKD (light blue), gp130 P216H (black), and gp130 V252G (purple).

**d** Zoomed-in view of D2D3 from structures in **c** showing distinct conformations of the βC'-C'E-βE modules in different mutants. Only one monomer in each mutant dimer is shown for better visibility.

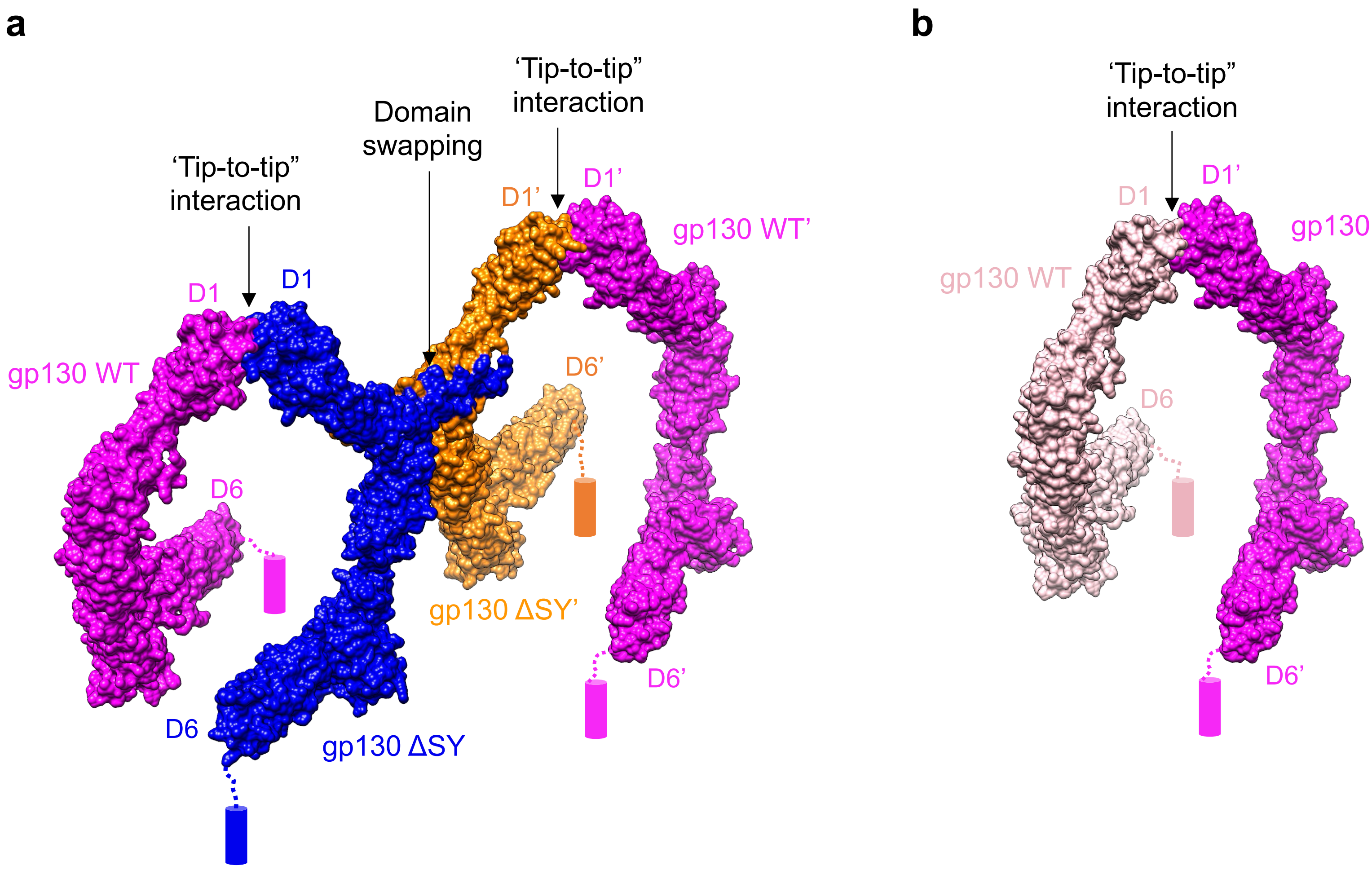


**Supplementary Figure 20. Hypothetical models of gp130 WT/ΔSY heterodimer (a) and pre-formed gp130 WT homodimer (b).** Cylinders represent TM domains.

**Supplementary Table 1. Cryo-EM data collection, processing, and refinement statistics**

| | ΔSY ECD dimer | D215G ECD dimer | V252G ECD dimer | ΔSY dimer | ΔSY dimer cluster | ΔSYΔD1 dimer |
|---|---|---|---|---|---|---|
| **Data collection/processing** | | | | | | |
| Magnification | 105,000 | 105,000 | 105,000 | 105,000 | 105,000 | 105,000 |
| Voltage (kV) | 300 | 300 | 300 | 300 | 300 | 300 |
| Electron exposure ($e^{-}/Å^{2}$) | 40 | 40 | 40 | 40 | 40 | 40 |
| Defocus range (μm) | -1.4 to -2.6 | -1.4 to -2.6 | -1.4 to -2.6 | -1.4 to -2.6 | -1.4 to -2.6 | -1.4 to -2.6 |
| Pixel size (Å) | 0.86 | 0.86 | 0.86 | 0.86 | 0.86 | 0.86 |
| Number of movies | 11,094 | 11,086 | 14,775 | 10,097 | 18,679 | 19,470 |
| Initial number of particles | 7,330,843 | 8,723,011 | 11,010,102 | 7,538,941 | 13,912,054 | 13,788,347 |
| Final selected particles | 291,116 | 220,794 | 229,453 | 242,322 | 69,303 | 63,329 |
| Symmetry imposed | C2 | C2 | C2 | C2 | C1 | C2 |
| Map resolution (Å) | 3.24 | 3.38 | 3.82 | 3.29 | 6.78 | 3.91 |
| FSC threshold | 0.143 | 0.143 | 0.143 | 0.143 | 0.143 | 0.143 |
| **Refinement** | | | | | | |
| Model composition | | | | | | |
| Non-hydrogen atoms | 15,802 | 15,874 | 15,876 | 15,802 | 31,604 | 14,268 |
| Protein residues | 2,014 | 2,024 | 2,024 | 2,014 | 4,028 | 1,820 |
| R.m.s. deviations | | | | | | |
| Bond lengths (Å) | 0.002 | 0.002 | 0.002 | 0.002 | 0.002 | 0.003 |
| Bond angles (°) | 0.494 | 0.495 | 0.500 | 0.610 | 0.633 | 0.801 |
| Validation | | | | | | |
| MolProbity score | 1.08 | 1.15 | 1.51 | 1.25 | 1.42 | 1.50 |
| Rotamer outliers (%) | 0.11 | 0.33 | 0.89 | 0.56 | 0.00 | 0.74 |
| Clash score | 2.92 | 3.58 | 4.69 | 4.85 | 7.75 | 9.57 |
| Ramachandran plot | | | | | | |
| Favored (%) | 98.60 | 98.61 | 96.12 | 98.50 | 98.40 | 98.34 |
| Allowed (%) | 1.40 | 1.39 | 3.88 | 1.50 | 1.60 | 1.55 |
| Disallowed (%) | 0.00 | 0.00 | 0.00 | 0.00 | 0.00 | 0.11 |
| **Deposition ID** | | | | | | |
| PDB | 9YJ5 | 9YJ6 | 9YJ7 | 9YJ8 | 9YJ9 | 9YJA |
| EMDB | 73011 | 73012 | 73013 | 73014 | 73015 | 73017 |

| | ΔSYΔ(D4-D6) dimer | ΔSYΔ(D4-D6) dimer cluster | ΔKD dimer | ΔEV dimer | P216H dimer | ΔAF dimer |
|---|---|---|---|---|---|---|
| **Data collection/processing** | | | | | | |
| Magnification | 105,000 | 105,000 | 105,000 | 105,000 | 105,000 | 105,000 |
| Voltage (kV) | 300 | 300 | 300 | 300 | 300 | 300 |
| Electron exposure ($e^{-}/Å^{2}$) | 40 | 40 | 40 | 40 | 40 | 40 |
| Defocus range (μm) | -1.4 to -2.6 | -1.4 to -2.6 | -1.4 to -2.6 | -1.4 to -2.6 | -1.4 to -2.6 | -1.4 to -2.6 |
| Pixel size (Å) | 0.84 | 0.84 | 0.84 | 0.84 | 0.84 | 0.84 |
| Number of movies | 14,147 | 14,147 | 10,906 | 10,134 | 19,118 | 19,193 |
| Initial number of particles | 10,688,683 | 10,688,683 | 5,937,687 | 5,329,929 | 12,904,732 | 8,723,011 |
| Final selected particles | 340,107 | 105,713 | 67,458 | 153,320 | 94,739 | 274,585 |
| Symmetry imposed | C2 | C1 | C2 | C2 | C2 | C2 |
| Map resolution (Å) | 3.04 | 6.35 | 3.36 | 4.02 | 4.09 | 3.62 |
| FSC threshold | 0.143 | 0.143 | 0.143 | 0.143 | 0.143 | 0.143 |
| **Refinement** | | | | | | |
| Model composition | | | | | | |
| Non-hydrogen atoms | 11,172 | 22,344 | 15,782 | 15,820 | 15,888 | 15,944 |
| Protein residues | 1,436 | 2,872 | 2,012 | 2,016 | 2,024 | 2,036 |
| R.m.s. deviations | | | | | | |
| Bond lengths (Å) | 0.002 | 0.004 | 0.002 | 0.002 | 0.002 | 0.003 |
| Bond angles (°) | 0.582 | 0.824 | 0.569 | 0.555 | 0.571 | 0.662 |
| Validation | | | | | | |
| MolProbity score | 1.30 | 1.59 | 1.56 | 1.53 | 1.75 | 1.52 |
| Rotamer outliers (%) | 0.16 | 0.00 | 0.78 | 0.00 | 0.00 | 0.77 |
| Clash score | 5.19 | 12.01 | 6.27 | 6.03 | 7.63 | 7.20 |
| Ramachandran plot | | | | | | |
| Favored (%) | 97.89 | 98.03 | 96.60 | 96.81 | 95.23 | 97.33 |
| Allowed (%) | 2.11 | 1.97 | 3.20 | 2.89 | 4.77 | 2.57 |
| Disallowed (%) | 0.00 | 0.00 | 0.20 | 0.30 | 0.00 | 0.10 |
| **Deposition ID** | | | | | | |
| PDB | 9YJB | 9YJC | 9YJD | 9YJE | 9YJF | 9YJG |
| EMDB | 73018 | 73019 | 73020 | 73021 | 73022 | 73023 |
